# Multifocal imaging for precise, label-free tracking of fast biological processes in 3D

**DOI:** 10.1101/2020.05.16.099390

**Authors:** Jan N. Hansen, An Gong, Dagmar Wachten, René Pascal, Alex Turpin, Jan F. Jikeli, U. Benjamin Kaupp, Luis Alvarez

## Abstract

Many biological processes happen on a nano-to millimeter scale and within milliseconds. Established methods such as confocal microscopy are suitable for precise 3D recordings but lack the temporal or spatial resolution to resolve fast 3D processes and require labeled samples. Multifocal imaging (MFI) allows high-speed 3D imaging but is limited by the compromise between high spatial resolution and large field-of-view (FOV), and the requirement for bright fluorescent labels. Here, we provide a new open-source 3D reconstruction algorithm for multifocal images that allows using MFI for fast, precise, label-free tracking spherical and filamentous structures in a large FOV and across a high depth. We characterize fluid flow and flagellar beating of human and sea urchin sperm with a high z-precision of 0.15 μm, in a large volume of 240 x 260 x 21 μm, and at high speed (500 Hz). The large sampling volume allowed to follow sperm trajectories while simultaneously recording their flagellar beat. Our MFI concept is cost-effective, can be easily implemented, and does not rely on object labeling, which renders it broadly applicable.

## Introduction

Life happens in three dimensions (3D). Organisms, cells, and sub-cellular compartments continuously undergo 3D movements. Many biological processes happen on the micrometer to millimeter scale within milliseconds: in one second, insects flap their wings 100 to 400 times^1,2^, microorganisms swim 0.5 to 50 body lengths^3–5^, cilia and flagella beat for up to 100 times^6,7^ and the cytoplasm of plants streams over a distance of 100 μm^8^. Although methods such as confocal and light-sheet microscopy allow precise 3D recordings, these techniques lack either the time or spatial resolution or are constrained to a small field-of-view (FOV) that is too narrow to follow fast biological processes in 3D. Rapid 3D movements can be studied by two complementary high-speed microscopy methods: digital holographic microscopy (DHM)^5,9–12^ and multifocal imaging (MFI)^13–16^.

DHM relies on the interference between two waves: a coherent reference wave and a wave resulting from the light scattered by the object. The 3D reconstruction of objects featuring both weak and strong scattering compartments, e.g., tail and head of sperm, is challenging because interference patterns of strong scattering objects conceal the patterns of weakly scattering objects. Additionally, DHM is very sensitive to noise and 3D reconstruction from DHM data requires extensive computations.

A simple alternative to DHM is MFI. MFI produces a 3D image stack of the specimen by recording different focal planes simultaneously. An MFI device, placed into the light path between microscope and camera, splits the light collected by the objective and projects multiple focal images of the sample onto distinct locations of the camera chip. However, this approach constrains the FOV^13^ and lowers the signal-to-noise ratio (SNR) when increasing the number of focal planes. Thus, state-of-the-art 3D tracking based on MFI requires bright fluorescent labels and is limited to either lower speeds, smaller sampled volumes, or lower precision^13,15–19^. Here, we develop dark-field-microscopy-based MFI for label-free high-speed imaging at a high SNR and across a large volume. We introduce a 3D reconstruction method for MFI that is based on inferring the z-position of an object based on its defocused image. This allowed tracking spheres and reconstructing filaments, such as flagella, at frequencies of 500 Hz, with sub-micrometer precision, in a large FOV of up to 240 x 260 μm, and across a large depth of up to 21 μm.

## Results

### Assembling a broadly applicable MFI setup

We aim to establish an MFI system based on dark-field microscopy that can deliver high-contrast images without sample labeling. Generally, five alternative approaches to simultaneously acquire different focal planes are exploited for imaging: using (1) an optical grating^13,18^, (2) a deformable mirror^20^, (3) a dual-objective microscope^21,22^, (4) different optical path lengths^14,16,23^, and (5) lenses of different focal power^15,17,19^. Several approaches suffer from some disadvantages: optical gratings are wavelength-dependent and have low optical efficiency. Deformable mirrors are bulky, expensive, and highly wavelength-dependent, which prohibits their use for low-cost compact applications. Finally, dual-objective microscopes are expensive and complex – e.g., they require two tube lenses and cameras – and are limited to two focal planes only and require fluorescent labels. Therefore, we aim for an MFI system that combines approaches (4) and (5) and thus, can be accommodated in different experimental set-ups. We use a multifocal adapter that splits the light coming from the microscope into four light paths that are projected to different areas of the camera chip (Supplementary Fig. 1a). By inserting four lenses of different focal power into the individual light paths, four different focal planes of the specimen are obtained (Supplementary Fig. 1b and 1d). Using only four focal planes minimizes the loss of SNR due to light splitting and keeps the system low in complexity. We established an alignment procedure based on recording images of a calibration grid (see Materials & Methods) to precisely overlay the four focal plane images (Supplementary Fig. 2) and to correct for subtle magnification differences between the four focal planes (Supplementary Table 1, 2). Of note, the images of the calibration grid did not show any apparent comatic aberrations, spherical aberrations, field curvature, astigmatism, or image distortions (Supplementary Fig. 1c, 2a). Using the thin-lens approximation, we can predict the defocusing of a pattern in the set-up based on the set of lenses and the magnification (Supplementary Fig. 1c and 1e). We assembled the multifocal adapter to study objects of submicrometer to millimeter size by changing the magnification (Supplementary Table 3). This flexibility enables studying fast-moving objects, ranging from subcellular organelles to whole animals.

### Extended depth-of-field imaging of fast-moving objects

Long-term imaging of a fast-moving object with high spatial and temporal resolution is a common, yet challenging goal in microscopy. Imaging at high resolution with objectives of high magnification and numerical aperture constrains the depth-of-field and the FOV, which increases the odds that the specimen exits the observation volume, thereby limiting the duration of image acquisition. This limitation can be overcome by combining images acquired at different depths using extended-depth-of-field (EDOF) algorithms^24^. However, this technique is not suited for fast-moving objects if different focal planes are acquired sequentially. MFI allows employing EDOF algorithms for fast-moving objects (Fig. 1). We combine MFI (Fig. 1a) and EDOF (Fig. 1b) (see Materials and Methods) to study living specimens over a broad range of sizes: grooming *Drosophila melanogaster* (Supplementary Movie 1), foraging *Hydra vulgaris* (Supplementary Movie 2), crawling *Amoeba proteus* (Supplementary Movie 3), and beating human sperm (Supplementary Movie 4). In each of the four different focal planes, distinct structural features appear sharply (Fig. 1c). Using EDOF algorithms, these regions are assembled into a single sharp image that reveals fine structural features, such as intracellular compartments (Supplementary Movie 3, Fig. 1b). The extended imaging depth allows tracking objects over long time periods and with high speed, even when the sample moves along the z-axis (Supplementary Movies 1-4). Of note, the EDOF method also reveals a depth map (Fig. 1c) that, in combination with MFI, allows coarse 3D tracking at high speed.

**Figure 1.**
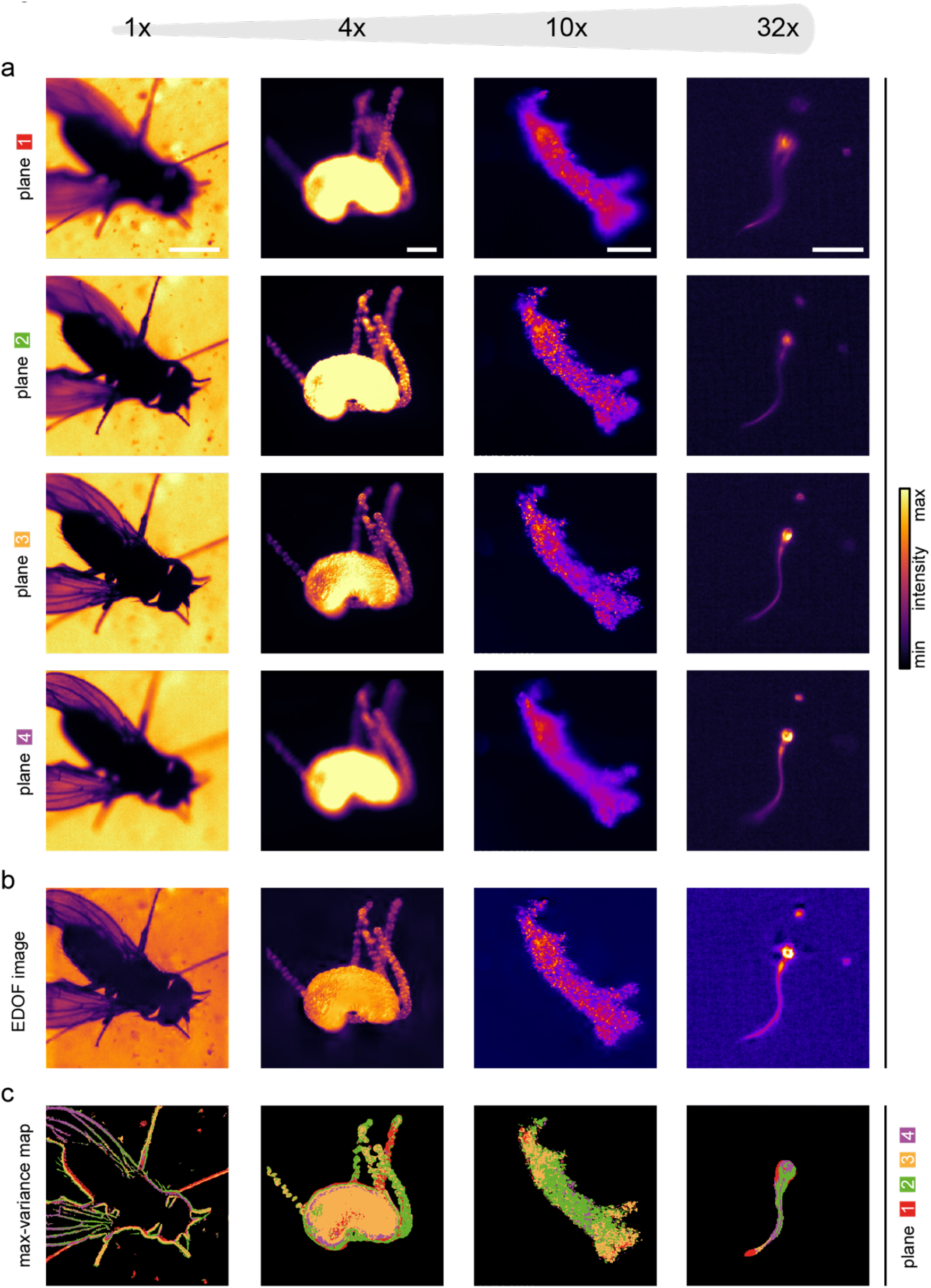
Multifocal imaging enables high-speed, extended depth-of-field visualization. **(a)** Multifocal images, acquired with the MFI system at various scales, demonstrate that MFI can be used to simultaneously image fast-moving objects at different depths. Left to right: grooming *Drosophila melanogaster* (1x; scale bar 1 mm, bright-field microscopy), foraging *Hydra vulgaris* (4x; scale bar 200 μm, dark-field microscopy), crawling *Amoeba proteus* (10x; scale bar 100 μm, dark-field microscopy), and swimming human sperm cell (32x; scale bar 20 μm, dark-field microscopy). **(b)** Extended depth-of-field (EDOF) images produced from multifocal images shown in (a). **(c)** Max-variance maps showing for specific pixel positions the planes (color-coded) wherein the specimen appeared most sharp (sharpness determined as pixel variance), revealing a coarse feature localization of the object in z.

### High-precision 3D tracking with MFI

MFI has been applied for 3D tracking^13,15–19^, but only with poor precision or a small sampling volume. The image of an object, i.e. the sharpness, size, and intensity of the image, depends on the object’s z-position relative to the focal plane. This dependance is used in “depth-from-defocus” methods to infer the z-position of an object based on the defocus of the object. Such methods have been applied to estimate the third dimension in physical optics^25–27^ and, for biological specimens, from 2D microscopy images^28–31^. To determine the z-position, a precise calibration of the relationship between the image of the object and the z-position is required. We combined the depth-from-defocus approach with MFI to achieve high-precision 3D tracking. We illustrate the power of this concept by reconstructing the 3D movement of spherical objects (latex beads) and filamentous structures (sperm flagella) using label-free imaging by dark-field microscopy and a multifocal adapter.

To calibrate the relationship between the image and the z-position of a spherical object, we acquired multifocal z-stacks of non-moving 500-nm latex beads by moving the objective with a piezo across a depth of 21 μm; the step size was 0.1 μm. The bead image underwent two characteristic changes as the objective was moved: the radius of the bead image increased (Fig. 2a), and, as a result, the image intensity decreased (Fig. 2b). Both features are z-dependent and can be used to determine the bead z-position. The z-position of a particle cannot be inferred unequivocally from a single plane. For any given bead radius, two possible z-positions exist (Fig. 2c). However, combining the information from multiple planes allows unequivocally localizing the particle in z (Fig. 2c) and, in addition, allows combining multiple measurements of the z-position in different planes by applying Bayesian inference probability^32^ or other statistical methods to improve the z-precision. We estimated the z-precision for different z-positions and for different focal planes based on the relationship between bead radius and bead z-position (see Materials & Methods) (Fig. 2d). The z-position near the focus is imprecise because the radius of the bead image hardly changes when the bead’s z-position is changed. By contrast, if the bead is slightly defocused, the relationship between the radius of the bead image and the z-position is steep, providing a better precision. When multiple focal planes are available to infer the z-position of a bead, the plane yielding the best precision can be selected to infer the z-position (Fig. 2e). Thereby, a better z-precision across the entire sampled volume is achieved. Increasing the number of planes allows increasing the range and z-precision (Fig. 2e). For the setup with four focal planes, we predicted a mean z-precision of 162 μm across a depth of 20 μm.

**Figure 2.**
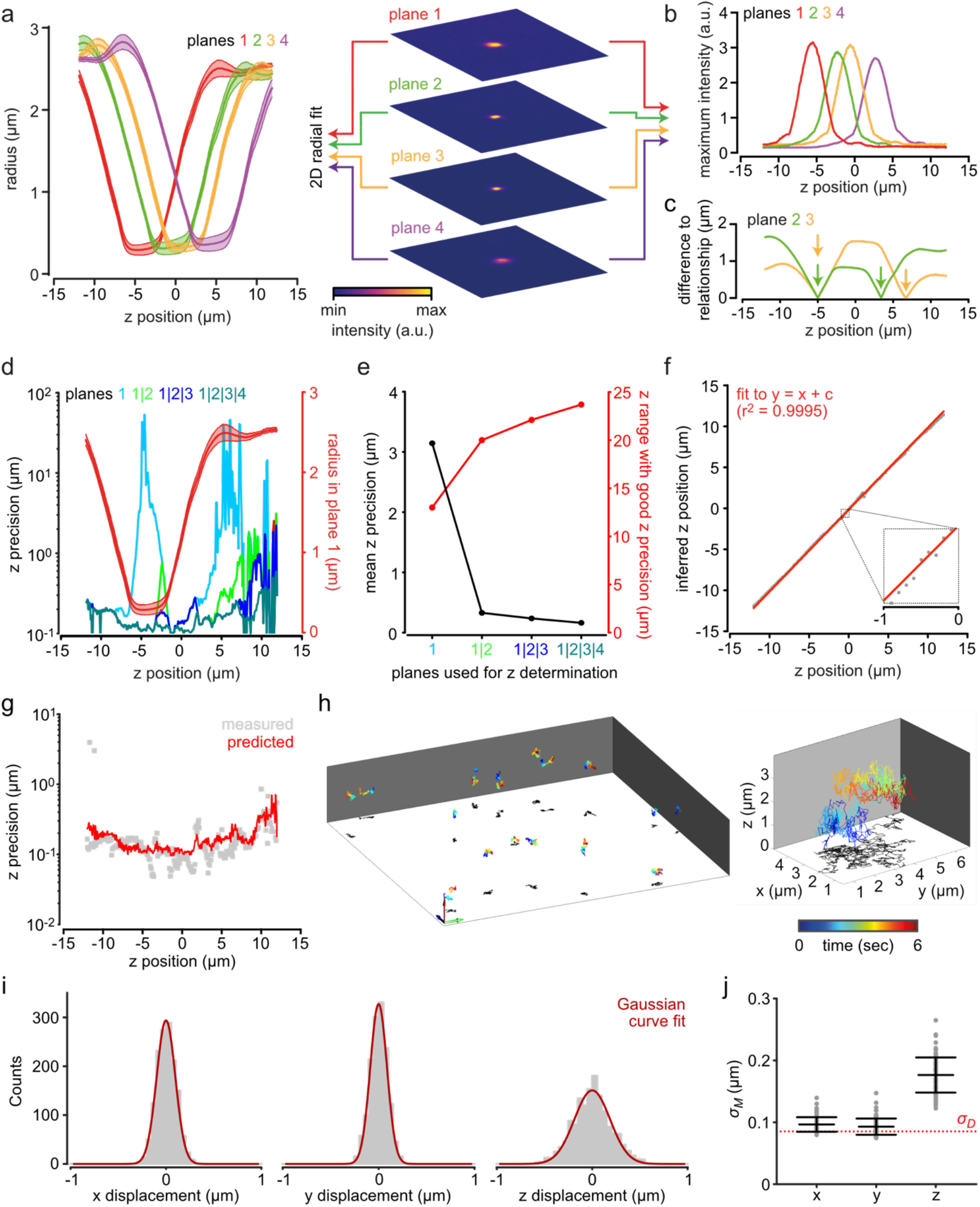
Localizing latex beads in z using four focal planes. **(a-c)** Characterizing the relationship between the image and the z-position of a latex bead (diameter 500 nm) in the MFI setup, equipped with a 20x objective (NA 0.5) and a 1.6x magnification changer. **(a)** Bead radius determined by a circle fit as a function of z-position; mean ± standard deviation of *n* = 7 beads (left). MF images of a latex bead at an exemplary z-position (right). **(b)** Maximum intensity of the bead image as a function of the bead’s z-position in the four focal planes for one exemplary bead. **(c)** Difference between the measured bead radius and the calibrated relationship between bead radius and z-position (from panel (a)) reveals two possible bead z-positions as minima (arrows) for each imaging plane. The overlay of the difference functions from two planes determines the bead’s z-position unequivocally. **(d)** Predicted z precision at different z-positions and **(e)** mean z-precision of all z-positions using plane 1 only, planes 1 and 2, planes 1 to 3, or planes 1 to 4. For comparison, the relationship between bead radius and z-position for plane 1 is overlaid in (d) (red). The range of z positions with a z precision better than 0.5 μm is overlaid in (e) (red). The z precision was predicted based on the calibrated relationship shown in (a) (see Materials and Methods). **(f)** z-position of a non-moving bead inferred from MF images based on the calibrated relationship between bead radius and z-position during modulation of the objective z-position with a piezo (step size: 0.1 μm). A linear curve with a unity slope was fit to the data. **(g)** z precision measured as the standard deviation of the residuals of linear curve fits with unity slope to the inferred z-positions during modulation of the objective z-position (gray), determined from *n* = 5 beads. The z precision predicted as in (d) was overlayed (red). **(h)** Representative 3D trajectories of freely diffusing latex beads, displaying a characteristic Brownian motion. Magnified view of an individual trajectory on the right. The z-positions of the beads were inferred by multifocal image analysis. Arrows indicate 10 μm. **(i)** Characterization of bead displacement between consecutive frames (*n* = 81 beads), demonstrating that the displacement of the bead is normally distributed in x, y, and z. **(j)** From the variance 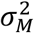 of the measured bead displacement and that predicted by diffusion 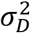 of the 500 nm beads (standard deviation shown as red dotted line) a localization precision of 45 nm, 37 nm, and 154 nm in *x, y*, and *z*, respectively, can be estimated (see materials and methods). Data points represent individual beads tracked for a mean duration of 3.3 s. *n* = 81 beads. Bars indicate mean ± standard deviation.

We next experimentally scrutinized the predicted z-precision. We recorded a new data set of multifocal z-stacks from immobilized latex beads using a piezoelectric-driven objective, measured the radius of the bead images in the different focal planes, and inferred the z-positions of the beads using the relationship between radius and z-position (Fig. 2 a) (see also Materials & Methods). Comparison of the inferred and piezo z-positions revealed a linear relationship (mean ± standard deviation of the slope: 0.98 ± 0.02, *n* = 5 beads). We determined the z-precision based on linear fits with a slope of unity (Fig. 2f); the predicted and experimentally determined z-precision closely matched across the entire depth (Fig. 2g).

To gauge the validity of our 3D-localization approach, we studied the stochastic Brownian motion of beads (*n* = 81) that were tracked for a mean duration of 3.3 s (Fig. 2h) and we determined the x, y, and z distributions of bead displacement (Fig. 2i). The distributions indicated that stochastic bead motion was isotropic and normal distributed over all spatial coordinates. From the distributions of bead displacement (Fig. 2i) we inferred the x, y, and z precision of our 3D-localization approach based on the assumption that the distributions represent the positional variations resulting from diffusion superimposed on the positional variations resulting from the measurement precision (see Materials & Methods, Fig. 2j). We obtained a z-precision of 45 nm, 37 nm, and 154 nm in x, y, and z, respectively. Of note, the z-precision agrees with the z-precision predicted and measured during the calibration procedure (Fig. 2g). We conclude that MFI is highly suitable for precise 3D tracking in a large volume.

### High-precision 3D reconstruction of the flagella beat with MFI

Eukaryotic cells such as sperm or green algae deploy motile, lash-like appendages protruding from the cell surface – called flagella. The 3D flagellar beat propels the cell on a 3D swimming path^29,33^. Because the beat frequency is high (up to 100 Hz)^6,7^, the precise reconstruction of flagellar beating at high temporal and spatial resolution is challenging. The fastest calibrated 3D reconstructions of human sperm achieved temporal resolutions of 100 frames per second (fps) with DHM^34^ and 180 fps using a piezo-driven objective^35^. This temporal resolution allows determining the beat frequency of human sperm that ranges, considering cell-to-cell variability, from 10 to 30 Hz^36^. However, this resolution is not sufficient to characterize higher harmonics of the beat^6,37^ or to track and characterize the first harmonic of fast-beating flagella, e.g. from *Anguilla* sperm (about 100 Hz)^7^. Detecting the second harmonic frequency (60 Hz) for a sperm beating at a fundamental frequency of 30 Hz will require, according to the Nyquist Shannon theorem, an acquisition rate of at least 120 fps. To determine the beat frequency of *Anguilla* sperm (about 100 Hz), an acquisition rate of at least 200 fps is required. Of note, the beat frequency of sperm is variable, rendering the determination of the flagellar beat frequency noisy. Thus, a temporal resolution of 120 fps would suffice only for regularly beating cells.

Recently, using holography, the flagellar beat of human sperm has been reconstructed with a high temporal resolution (2,000 fps)^38^. However, the small spectral power compared to the first harmonic and the low number of beat cycles sampled during the short recording time (0.5 s) did not allow for a clear detection of the second harmonic. Moreover, the tracking precision was not evaluated and the sampled volume was very small (ca. 30 x 45 x 16 μm)^38^ compared to the flagellum length (50 μm), which compromises deriving the relation between flagellar beat and trajectory.

We aimed to establish MFI for reconstructing the 3D flagellar beat of human sperm at high speed (500 Hz), with high precision, and in a large FOV that allows tracking both the flagellar beat and trajectory over a long time.

We used a magnification of only 32x, which decreased the optical resolution compared to previous methods, employing 100x magnification^17,38^. However, this yielded a 58-times larger sampled volume of ca. 240 x 260 x 20 μm compared to a DHM-based method (30 x 45 x 16 μm)^38^ and an 80-times larger sampled volume compared to an MFI-based method (ca. 80 x 35 x 5.6 μm)^17^. We compensated for the lower optical resolution by establishing a precise reconstruction method similar to that for bead tracking.

To characterize the relationship between image and z-position of flagella, we acquired multifocal z-stacks of immotile sperm using the piezo-driven objective. Similar to the latex beads, the flagellum image displayed a characteristic widening (Fig. 3a) and an intensity decrease (Fig. 3b) when imaged out of focus.

**Figure 3.**
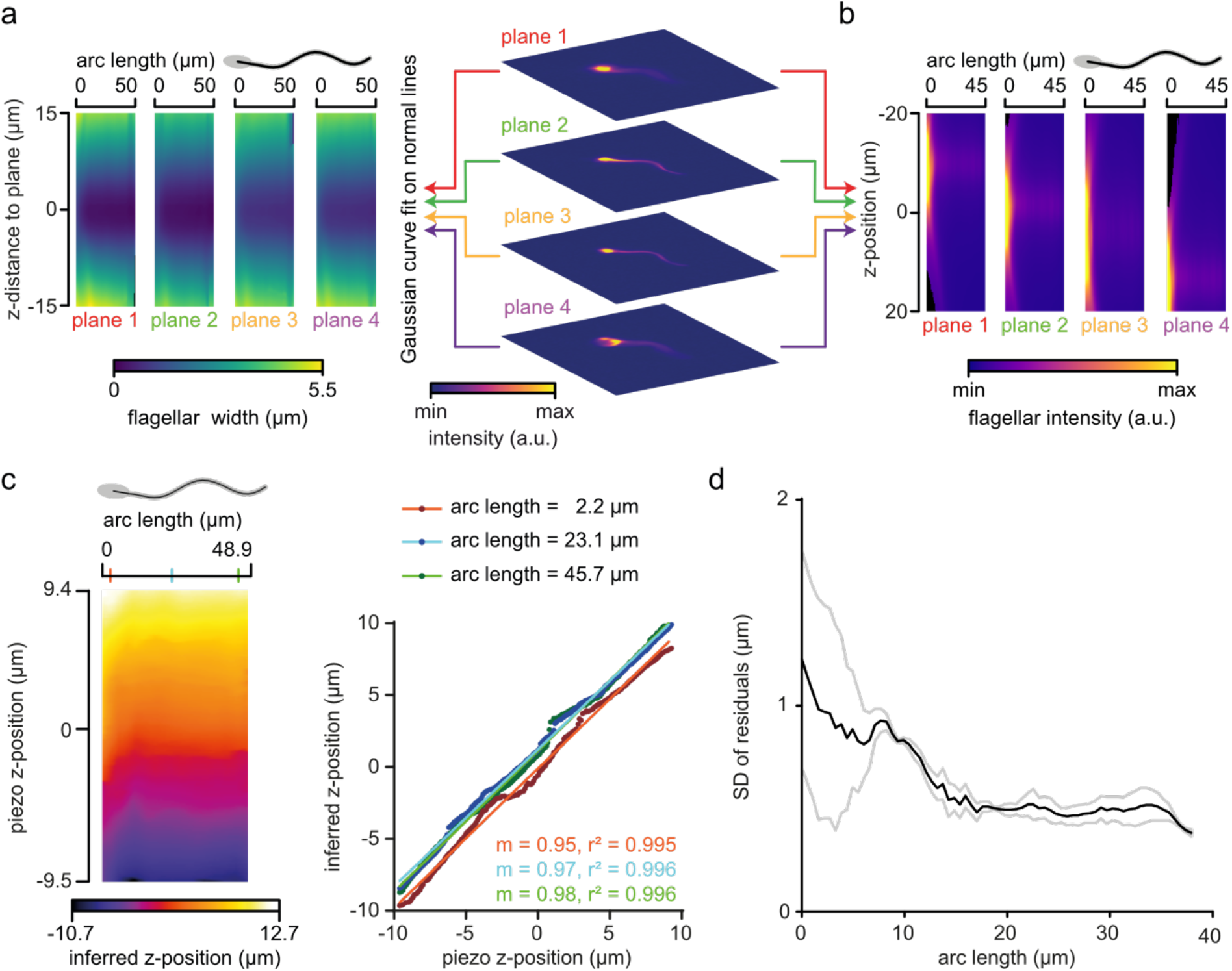
Reconstructing flagellar z-positions of human sperm using four focal planes. **(a-b)** Characterizing the relationship between image and z-position of human sperm flagella in the MFI setup. **(a)** Flagellar width (color-coded), determined by a Gaussian curve fit on a normal to the flagellum, as a function of the flagellar position (arc length) and the z-distance to the respective plane (mean image of *n* = 12 sperm from five different donors). **(b)** Maximum flagellar intensity (color-coded) as a function of the flagellar position (arc length) and the z-position relative to the four planes. **(c)** z-position (color-coded) along the flagellum (arc length) of an immotile human sperm cell inferred from MF images based on the calibrated relationship between the flagellar width, position on the flagellum, and z-distance to the respective plane during modulation of the objective z-position with a piezo (steps 0.1 μm; left). Colored ticks on the arc-length axis mark flagellar positions that are further analyzed by a linear curve fit (right), revealing a linear relationship between the objective z-position and the z-position determined by MF image analysis (m: slope). **(d)** Standard deviation (SD) of the residuals of linear curve fits to data as exemplified in (c). Mean ± standard deviation of *n* = 3 sperm of different donors.

We refined SpermQ, a software for analyzing motile cilia in 2D^39^, to incorporate the analysis of multifocal images (SpermQ-MF); SpermQ-MF determines flagellar z-positions based on the calibrated relationship between flagellar width, position along the flagellum, and z-distance to the respective plane. In each plane of multifocal images, SpermQ reconstructs the flagellum in 2D (x and y coordinates) with a good precision of about 0.05 μm for the focused and about 0.1 μm for the defocused flagellum (Supplementary Fig. 3). To determine the z-precision, we inferred the z-position of an immotile sperm using SpermQ-MF in multifocal z-stacks recorded with the piezo-driven objective (Fig. 3c). Inferred and piezo z-positions showed a linear relationship with a slope of unity (Fig. 3c). From linear fits to the relationships of inferred and piezo z-position at different arc length positions (*n*= 3 sperm) (Fig. 3c), we determined a z-precision of ~1 μm at the head and midpiece and 0.6 μm at the principal piece (Fig. 3d).

We next tested whether our approach is applicable to flagella of other species by applying the same procedure to sperm from the sea urchin *Arbacia punctulata* (Supplementary Fig. 4). Here, fits to determine the flagellar width at the neck region were compromised by the highly flexible neck and the high-intensity contrast between head and flagellum in sea urchin sperm. Thus, the z-precision at the flagellar neck region (arc length < 10 μm) was twofold less for sea urchin sperm (~2 μm, Supplementary Fig. 4c-d) than for human sperm (~1 μm, Fig. 3d). In the more distal part of the flagellum (arc lengths > 10 μm), sea urchin sperm were reconstructed with a better z-precision (~0.4 μm, Supplementary Fig. 4d) compared to human sperm (~0.6 μm, Fig. 3d).

In conclusion, our approach provides high temporal resolution (500 Hz), a large FOV of ca. 240 x 260 x 20 μm, and allows flagellar 3D reconstruction with a z-precision of down to ~0.4 μm, whereas for other methods the z-precision has not been reported^17,38^.

### 3D reconstruction of the flagellar beat with MFI

We next studied and compared the flagellar beat of free-swimming sperm from human and sea urchin. Using the calibrated MFI setup and SpermQ-MF, we recorded the 3D flagellar beat of *n*= 6 human and *n* = 10 sea urchin sperm swimming in a 150-μm deep chamber. For human sperm, the tracking duration was limited by the FOV to about 2-3 s, i.e. the time it takes a sperm cell crossing the FOV (Fig. 4a, Supplementary Movie 5). Due to the precise localization of the flagellum, we resolved the 3D flagellar beat in detail (Fig. 4b, c). The kymographs revealed a flagellar beat wave that propagates in 3D along the flagellum (Fig. 4b).

**Figure 4.**
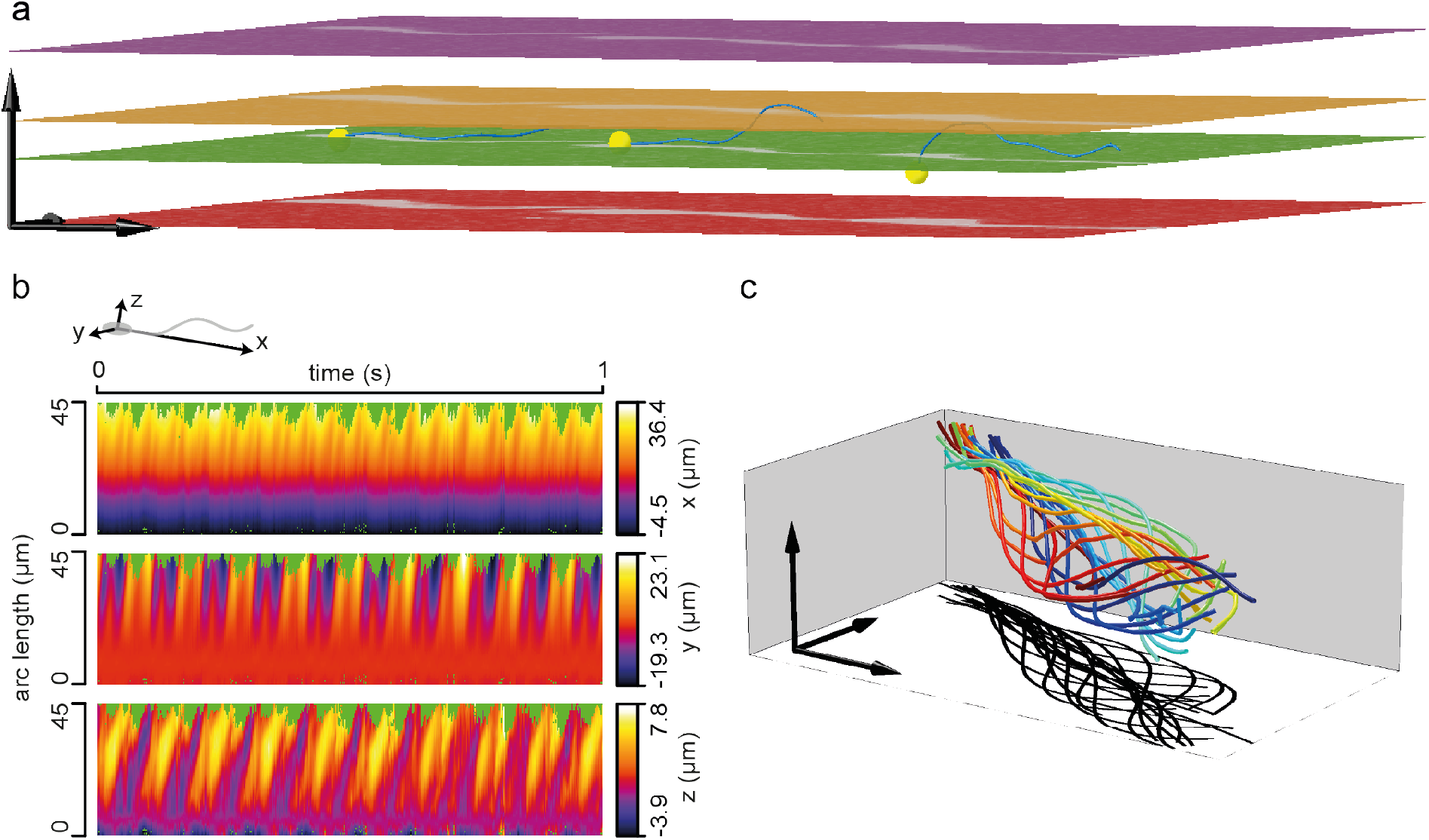
3D reconstruction of flagella from a swimming human sperm cell. **(a)** 3D visualization of the four planes (depicted in different colors) acquired by MFI and the flagellum reconstructed using SpermQ-MF and the calibrated relationship between flagellar width, position along the flagellum, and z-distance to the respective plane (Fig. 3a). Overlay of three exemplary time-points. Flagella indicated in blue. Positions of sperm heads indicated as yellow spheres. Arrows indicate 20 μm. **(b)** Kymographic representation of flagellar 3D coordinates (color-coded) in a reference system defined by the head-midpiece axis (see sketch on top). For better visualization, only the first second was plotted (reconstructed time span: 2.2 s). **(c)** 3D visualization of one beat cycle (time is color-coded). Arrows indicate 10 μm. Shadow indicates a projection of the flagellar beat to the xy-plane.

Making use of the large FOV of our MFI system, we recorded the flagellar beat and swimming path over a time span that allows relating beat pattern and trajectory (Fig. 5, Supplementary Fig. 5-8). Trajectories of human sperm varied between cells and over time for single cells (Fig. 5a, Supplementary Fig. 6). For example, one sperm cell changed from a slightly curved to a straight trajectory (Supplementary Fig. 6a); another sperm changed swimming directions multiple times (Supplementary Fig. 6e). By contrast, sea urchin sperm swam on a regular, circular swimming path based on a curved beat envelope (Fig. 5b, Supplementary Fig. 7-8). Sea urchin sperm can spontaneously depart from circular swimming to produce a stereotypical motor response known as “turn-and-run”^40^. Making use of the large FOV of our method, we were able to record a sperm during such a motor response and correlate the swimming trajectory and the flagellar beat (Supplementary Fig. 9). The flagellar beat transitioned from an asymmetric beat during the “turn” to a symmetric beat during the “run” (Supplementary Fig. 9).

**Figure 5.**
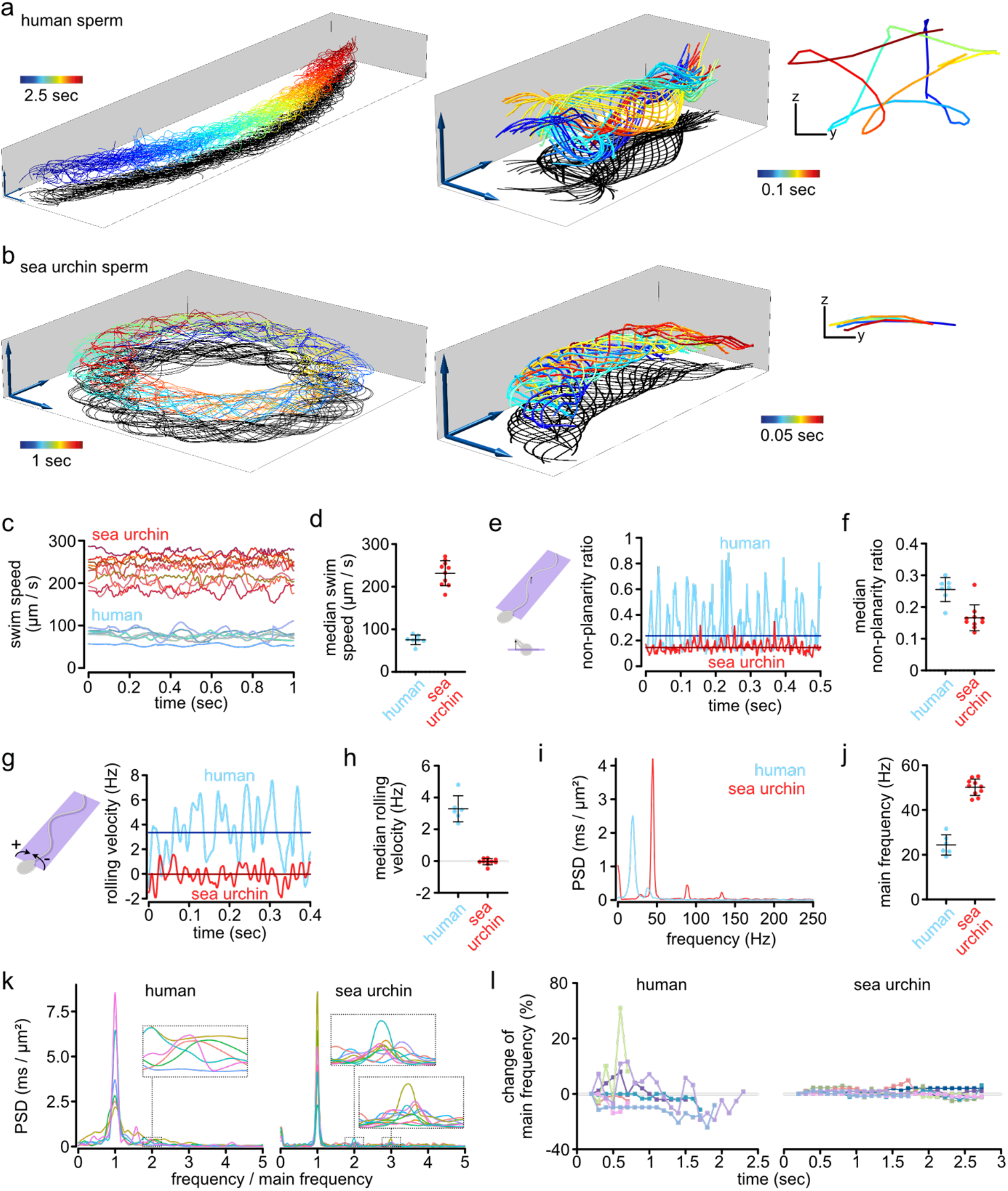
Relating flagellar beat pattern and swimming trajectory of human and sea urchin sperm. **(a-b)** 3D-tracked flagella from exemplary free swimming, **(a)** human and **(b)** sea urchin sperm. Views for visualizing the trajectory (left, only every 4^th^ frame plotted for better visualization), the flagellar beat (middle), or sperm rolling (right, a Y-Z-projection of a flagellar point at arc length 20 μm, view in swimming direction). Bars and arrows indicate 10 μm. Additional tracked sperm are displayed in Supplementary Fig. 6-8) **(c-l)** Quantification of the swimming and flagellar beating of all tracked sperm (*n* = 6 human sperm from two different donors and *n* = 10 sea urchin sperm). **(c-d)** 3D Swim speed, calculated as described in Supplementary Fig. 5. Each line (c) or point (d) corresponds to one tracked sperm cell. **(e-f)** Non-planarity ratio. Exemplary time courses (e) and median over time (f) of the non-planarity ratio. The non-planarity ratio is determined as the ratio of the two minor Eigenvectors of the flagellar inertial ellipsoid^29^. Dark lines in (e) show the median over time for the entire time course. Each data-point in (f) corresponds to the single-sperm median. **(g-h)** Rolling velocity. Exemplary time courses (g) and median over time (h) of the rolling velocity. Positive values indicate clockwise rotation when viewing the sperm from the head to the tail. Dark lines show the median obtained from the whole time series. Each data point in (h) corresponds to the singlesperm median. **(i-l)** Frequency spectrum of the flagellar beat in the distal flagellum. **(i)** Frequency spectra for an exemplary human and an exemplary sea urchin sperm (determined at arc length 33 μm and 34 μm, respectively). **(j)** Frequency of the highest peak in the frequency spectra of all human and sea urchin sperm analysed (determined at arc length 33 μm and 34 μm, respectively). Individual data points represent individual sperm. **(k)** Frequency spectra of all human and sea urchin sperm analysed after normalization of the frequency axis in each spectrum to the frequency of the highest peak (spectra determined at arc length 33 μm). **(l)** Frequency of the highest peak in the frequency spectrum as a function of the time (mean frequency of arc lengths 20-30 μm shown). Each data point represents the analysis of the time span from 0.2 s before to 0.2 s after the indicated time point. Bars indicate mean ± standard deviation.

We further quantified the 3D trajectory and the 3D flagellar beat (Fig. 5c-l). Human sperm swam at lower speed than sea urchin sperm (Fig. 5c-d). Additionally, the flagellar shape of human sperm was characterized by a large out-of-plane component (Fig. 5a) while the flagellar shape of sea urchin sperm was rather planar (Fig 5b). We quantified the out-of-plane component using the non-planarity ratio^29^ (see also Materials and Methods). This ratio ranges from zero to one. A ratio of zero indicates a perfectly planar beat, whereas a ratio of unity indicates a flagellar beat that has no preferential plane. The ratios largely fluctuated over time, especially for human sperm (Fig. 5e): The mean standard deviation over time of the non-planarity ratio was 0.17 ± 0.02 (mean ± standard deviation of *n* = 6 sperm) in human sperm and 0.08 ± 0.03 (mean ± standard deviation of *n* = 10 sperm) in sea urchin sperm. To exclude any bias by large fluctuations, we determined the median of the ratio over time as a reference for the nonplanarity per sperm cell (Fig. 5e, f). The median non-planarity ratio was 0.26 ± 0.04 (mean ± standard deviation of *n* = 6 sperm) in human sperm and 0.17 ± 0.04 (mean ± standard deviation of *n* = 10 sperm) in sea urchin sperm (Fig. 5f), confirming a more planar beat of sea urchin compared to human sperm. In addition, the beat plane in human sperm rolled around the longitudinal sperm axis (Supplementary Movie 6), whereas no rolling was detected for sea urchin sperm (Fig. 5g, h). Our measurements of the non-planarity ratio (0.26) and the rolling velocity (3.3 Hz, representing 20.7 rad/s) agree with a previous analysis using phase-contrast microscopy^29^.

Sperm steering has been related to asymmetric flagellar beat patterns that can be described by superposition of three components: (1) a mean curvature component C_0_, (2) the curvature amplitude *C*_1_ of the bending wave oscillating with the fundamental beat frequency, and (3) the curvature amplitude of the second harmonic C_2_^6,37^ To visualize higher harmonic components of the flagellar beat, we determined the frequency spectrum of the signed flagellar 3D curvatures (see Materials and Methods) for human and sea urchin sperm (Fig. 5 i-j). The principal beat frequency was 24.4 ± 4.5 Hz for human and 50.2 ± 3.6 Hz (mean± standard deviation) for sea urchin sperm (Fig. 5 f), in line with other reports^12,41,42^. We observed higher harmonics in the spectrum of human and sea urchin sperm (Fig. 5i). The curvature components *C*_0_ and *C*_1_ were much more prominent in sea urchin compared to human sperm, whereas the second harmonic component *C*_2_ was only marginally different between sea urchin and human (Supplementary Fig. 10). We determined a second harmonic component of *C*_2_ = 0.006 ± 0.002 μm^-1^ (mean ± standard deviation) for human sperm. Notably, the widths of the frequency peaks were narrower for sea urchin than for human sperm (Fig. 5i, k), indicating that the beat frequency of human sperm is less constant than the beat frequency of sea urchin sperm. To confirm this observation, we analyzed the beat frequency of human and sea urchin sperm at different recording times, which revealed large variations in the beat frequency of human sperm over time but only small variations for sea urchin sperm (Fig. 5l). Such variations have not been reported in previous analysis, where a constant beat pattern with sharp frequency peaks was measured^6,37^. However, these measurements relied on tethering sperm at their head to establish sufficiently long observations for resolving the power-spectrum. When tethered, human sperm are prevented from rolling – they are forced into a stable condition and their beat plane is aligned with the glass surface by hydrodynamic interactions^43^. In contrast, our technique, due to precise tracking in a large FOV, allows recording freely-swimming human sperm under unconstraint conditions and over a sufficiently long time to detect variations in the flagellar beat.

### 3D particle imaging velocimetry to reconstruct fluid flow around a human sperm cell

The behaviour of ciliated cells and the underlying physics is of great interest for cell biology and biotechnological applications. We are only beginning to understand how sperm manage to navigate in a highly complex environment like the female genital tract^44^ or how motile cilia synchronize their beat to produce fluid flow^45^. Key to understanding these phenomena are the hydrodynamic interactions of motile cilia with each other and their aqueous environment^4,45^. The fluid flow resulting from flagellar beating has been studied experimentally and theoretically. Experimental studies resolved the 2D flow pattern for different microorganisms such as *Giardia lamblia*^46^, *Chlamydomonas reinhardtii*, and *Volvox*^47^. The 2D flow profile around human sperm has been inferred from the flagellar beat^48^, and the 3D profile from numeric simulations of the flagellar beat of sea urchin sperm near walls^43^. However, the predicted flow profiles have not been tested experimentally.

To visualize the 3D flow around human sperm with our MFI technique, we used latex beads whose relationship between radius and z-position had been previously calibrated. To follow the direction of the bead movement relative to the sperm flagellum, we tethered the sperm head to the glass surface.

Bead tracking revealed long-range hydrodynamic interactions between sperm and beads (Fig. 6a, Fig. 6b, Supplementary Movie 7). At large distances from the sperm cell, beads displayed a random Brownian motion (Fig. 6c). Near the cell, beads displayed a Brownian motion with a drift towards the sperm head (Fig. 6d) and away from the flagellar tip (Fig. 6e). Near the flagellum, beads described a fast 3D spiral-like movement resulting from the 3D flagellar beat (Fig. 6d-e).

**Figure 6.**
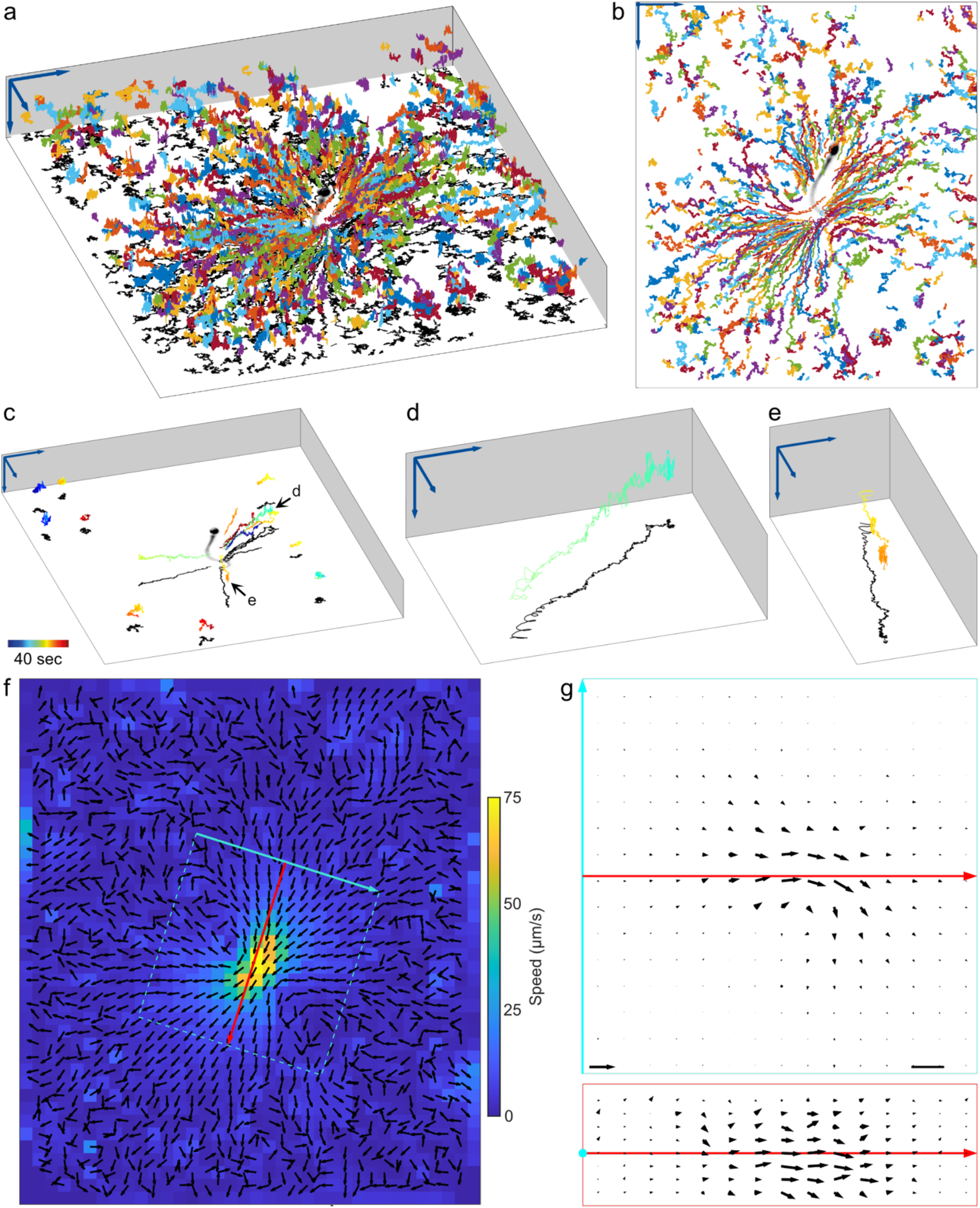
3D particle imaging velocimetry around a sperm cell. **(a-b)** Exemplary 3D trajectories of latex beads flowing around a human sperm cell tethered to the cover glass at the head. Each trajectory is depicted in a different colour. Arrows indicate 20 μm. Perspective (a) and top (b) views are shown. The bead trajectories have been projected onto an image of the tracked sperm cell (intensity inverted for better visualization, beads removed by image processing). Arrows indicate 20 μm. **(c)** Trajectories (color-coded by time) of few exemplary beads from (b). Beads that are remote from the flagellum show Brownian motion; close to the sperm flagellum the beads display a 3D spiralling motion. Time color-coded as indicated. Black arrows mark trajectories magnified in (d) and (e). The other trajectories shown are magnified in Supplementary Fig. 11. **(d)** Trajectory of a bead that is attracted to the sperm cell. **(e)** Trajectory of a bead that moves away from the flagellum. **(f)** Averaging z-projection of the 3D flow profile obtained from the bead motion, for which exemplary trajectories are shown in (a) to (e). The cyan and red arrows indicate a new coordinate system, which is used as a reference to show the 3D flow profile in (g). Black arrows are normalized. Flow speed is color-coded. **(g)** Flow profile sections in the vicinity of the sperm flagellum, extracted from the 3D flow at the positions marked with cyan and red lines / rectangles in (f). Cross-section (bottom) and top view (top) are shown. Scaler bar: 5 μm. Arrow on bottom left indicates 125 μm / s.

We computed the 3D flow field around a human sperm based on a time-lapse recording of 40 s at 500 Hz, revealing 2,920,731 bead positions with a predicted median z-precision of 0.34 μm (prediction based on the approach shown in Fig. 2d; median used to exclude outliers in this calculation). The flow field shows that the fluid moves from a large distance towards the sperm cell and converges at the flagellum, where the flow speed steeply increases (Fig. 6f, g). This agrees with the flow profiles predicted by theory^4,43,48^. We observed a flow speed of up to 125 μm s^-1^ directly at the flagellum (Fig. 6g). Ishimoto et al. predict for the flow speed close to the flagellum of human sperm a value of 0.1 *L/T*, where *L* corresponds to the length of the flagellum and *T* to the duration of one beat cycle^48^. For a beat frequency of 25 Hz and a flagellar length of 50 μm, this corresponds to a flow speed of 125 μm s^-1^, in perfect agreement with our experimental results. For human sperm, fluid flow at the flagellum has been only measured in 2D, revealing a flow speed of up to 45 μm s^-1^ in low-viscosity medium^49^. The flow speed derived from projecting our data into 2D by averaging the speeds across z are in a similar range (Fig. 6f). Taken together, our 3D analysis confirms predictions based on mathematical modelling of the 3D fluid flow around sperm and highlights the importance of measuring flow in 3D to achieve accurate estimates of the flow speed. Finally, we unravel spiral flow patterns close to the sperm cell.

## Discussion

State-of-the-art MFI systems that employ four focal planes have been limited to a depth < 6 μm and use high-magnification objectives (≥ 60x) to resolve the specimen, thereby constraining the FOV (≤ 80 x 35 μm)^15,17^ Although such a small FOV allows reconstructing the beat period of a freely-swimming human sperm (flagellum length: 50 μm), it is impractical to relate the beat pattern to the swimming trajectory because cells quickly leave the FOV.

Here, we present a new, depth-from-defocus-based 3D reconstruction algorithm that allows to computationally achieve an unprecedented z-precision in 4-plane multifocal images as high as 0.15 μm. Thereby, we are able to use a lower magnification (32x) and a wider plane spacing (planes spanned 8.8 μm). Using this algorithm, we precisely reconstruct beads and flagella across a 21-μm depth, a FOV of up to 240 x 260 μm, and at a high volume-sampling rate (500 Hz). This represents a 3.5-fold increase of depth, a 22-fold increase of FOV, and a 2.5-fold increase of sampling speed compared to other MFI systems^15,17^. The sufficiently large FOV allows relating swimming trajectories and beat pattern of sperm without compromising precision and speed.

For sperm, depth-from-defocus methods have been applied to roughly estimate the z-position in single-plane microscopy, without considering the z-precision^29,39^. We demonstrate that the z-precision inferred from a single plane is not equal for different z-positions and poor when the object is located near focus. We have surmounted these shortcomings by using multiple focal planes to derive the z-position and show that this allows unambiguous determination of the z-position, extensive improvement of the z-precision, approximation of the z-precision over a large depth, and significant increase in depth compared to using only one focal plane. Of note, these improvements depend on a well-designed MFI setup. To establish a good tracking precision over a high depth range, the interplane distance needs to be chosen based on the relationship between object width and z-position for an individual plane image.

Our MFI system represents a cost-effective 3D imaging device that does not rely on any special equipment and uses a commercially available adapter that is compatible with most microscopes. The system is flexible: it allows customizing the focal distance between image planes using different adapter lenses and it can be adapted to object sizes ranging from nano-to millimeter using different objectives. Thus, our MFI system is affordable and accessible to a large scientific community. To allow broad accessibility and applicability of our computational methods, we have implemented the whole image analysis pipelines presented in this study as plugins for the free, open-source software ImageJ (https://github.com/hansenjn/MultifocalImaging-AnalysisToolbox). Our software features a user interface and does not require any programming knowledge. Thus, we envision a broad applicability of these methods.

Alternative methods to derive depth information from a single-plane image use chromatic and spherical aberrations or astigmatism^50–56^. Like our method, these methods cannot resolve the z-position of objects that are stacked above each other in z or that are very close. Of note, to avoid any bias from this limitation, we implemented an algorithm that excludes objects with low-quality fits (Materials and Methods). Although deriving the depth via aberrations or astigmatism allows deciding whether an object lies before or behind the focal plane from one image only, these methods suffer from other disadvantages. First, using chromatic aberrations for depth estimates, multiple colors have to be acquired at the same time, and the object shape in different colors is compared to determine the z-position. In contrast to depth-from-defocus methods, this is not applicable to fluorescent samples. Moreover, it will lower the SNR and image contrast similarly to splitting light into multiple paths in our MFI setup. Second, astigmatic imaging to derive depth information has been only established for single-molecule imaging and spherical particles^53,55–57^. 3D tracking of flagella by astigmatic imaging or spherical aberrations is challenging, because the flagellar orientation in the focal plane and, for spherical aberrations also the flagellar x, y-position in the FOV, modulate the flagellar image acquired by these techniques. Thus, extensive calibration of additional variables that take these modulations into account would be required.

Previous studies of sperm’s flagellar 3D beat used light microscopy with a piezo-driven objective or digital holographic microscopy (DHM)^34,38,58–61^. The piezo-driven method allows recording up to 180 volumes s^-1^^35,61^, which represents a third of the sampling speed achieved by our MFI method. Of note, the piezo-driven method sequentially acquires multiple focal planes and, thus, requires high-speed high-sensitivity cameras that can record at high frame rates. For example, previous reconstructions of the flagellar beat using the piezo-driven method required acquisition of 8,000 images s^-1^^35,61^. Cameras achieving such acquisition rates are very expensive, restricting the availability of this technique to few labs. DHM can provide the same temporal resolution as MFI and allows high-precision 3D particle imaging velocimetry (PIV) achieving a precision down to 10 nm for simple spherical objects^62–64^. However, such precision is not achievable for biological samples: for cells, precisions of about 0.5 μm have been reported^65,66^. There are more factors that limit DHM. First, DHM is challenging when imaging strong- and weak-scattering objects at the same time; therefore, by contrast to MFI, DHM would not be capable of reconstructing a “small” sperm cell close to a “large” egg. Similarly, tracking the fluid flow around sperm may be challenging with DHM. Second, DHM is not applicable to fluorescent objects, whereas, using different labels, MFI could reveal different objects or processes simultaneously^13,15–19^. For example, applying our 3D reconstruction method to multicolor MFI allows to simultaneously record the 3D flagellar beat and fluorescent indicators for subcellular signaling. Third, DHM requires complex and extensive computational methods, whereas MFI does not. A state-of-the-art DHM-based reconstruction method for flagella featured a high temporal resolution (2,000 volumes s^-1^) but a small sampled volume of ca. 30 x 45 x 16 μm; the precision of the method was not reported^38^. Our technique for 3D reconstruction of human sperm provides a unique combination of a large sampled volume (ca. 240 x 260 x 20 μm), precision (≥ 0.4 μm), and high speed (500 Hz), which is superior to currently available alternative methods. Therefore, we could record higher harmonic frequencies in the flagellar beat of swimming sperm and follow the sperm trajectory upon a change in the flagellar beat pattern.

Present techniques allow studying the fluid flow surrounding flagellated microorganisms in 2D^46–48^. The 3D profile has previously been estimated by numerical simulations of the 3D flagellar beat of sperm^43^. Applying our technique for 3D particle imaging velocimetry allowed for the first time to record in 3D the fluid flow around a motile cilium, i.e. the human sperm flagellum. We reveal 3D spiral-shaped flow patterns around sperm. Moreover, we show that fluid flow transports molecules from afar to the flagellum. The flagellum acts as a propeller and a rudder, whose function is regulated by extracellular cues that act on intracellular signalling pathways^4^. Thus, it will be important to further investigate how the spiral-shaped flow pattern and the transport of fluid to the flagellum impact the sensing of chemical or mechanical cues by sperm during chemotactic or rheotactic navigation, respectively. Combining MFI and fluorescence microscopy^13,15–17,21–23,67^ allows simultaneously 3D tracking of the sperm flagellum and performing 3D particle imaging velocimetry using different fluorescent labels. This will reveal how a flow profile is generated by the 3D beat and how fluid flow varies in time as the beat progresses^68^.

Our method does not rely on a particular microscope configuration, which ensures broad applicability. We envisage three additional applications.

First, although we focused here on label-free samples using dark-field microscopy, our method is directly adaptable to fluorescently-labelled samples. Dark-field microscopy with a standard condenser, as applied here, limits the application to moderate NA objectives and consequentially also limits the optical resolution. However, using oil-immersion dark-field condensers or fluorescence-based MFI, high NA objectives could be applied, improving optical resolution.

Second, our MFI-based filament reconstruction method could be applied to larger objects, for instance to study rodent whisking during active vibrissal sensing^69^.

Third, we show that the concept of combining depth-from-defocus with multiple focal planes significantly improves depth estimates. Depth estimates are not only relevant for high-speed microscopy but also for research in 3D object detection by computer vision, including autonomous driving.

## Material and Methods

### Species

*Amoeba proteus* and *Hydra vulgaris* were purchased freshly before the experiments (Lebendkulturen Helbig, Prien am Chiemsee, Germany). *Drosophila melanogaster* were adults (> 2 days old). Human semen samples were donated by healthy adult males with their prior written consent and the approval of the ethics committee of the University of Bonn (042/17). Human sperm cells were purified by a “swim-up” procedure^70^ using human tubular fluid (HTF) (in mM: 97.8 NaCl, 4.69 KCl, 0.2 MgSO_4_, 0.37 KH_2_PO_4_, 2.04 CaCl_2_, 0.33 Na-pyruvate, 21.4 lactic acid, 2.78 glucose, 21 HEPES, and 25 NaHCO_3_ adjusted to pH 7.3–7.4 with NaOH). Sea urchin sperm from *Arbacia punctulata* were obtained from the Marine Resource Center at the Marine Biological Laboratory in Woods Hole. 200 μl of 0.5-M KCl were injected in the body cavity of the animals. Sperm were collected using a Pasteur pipette and stored on ice. Sperm suspensions were prepared in artificial seawater (ASW) at pH 7.8, which contained (in mM): 423 NaCl, 9 KCl, 9.27 CaCl_2_, 22.94 MgCl_2_, 25.5 MgSO_4_, 0.1 EDTA, and 10 HEPES.

### Imaging

All images except those of *D. melanogaster* were acquired using an inverted microscope (IX71; Olympus, Japan) equipped with a dark-field condenser and a piezo (P-725.xDD PIFOC; Physik Instrumente, Germany) to adjust the axial position of the objective. For MFI, the multi-channel imaging device (QV-2, Photometrics, USA) was installed in front of a high-speed camera (PCO Dimax, Germany). Different objectives were used: 20x (UPLFLN, NA 0.5; Olympus, Japan), 10x (UPlanSapo, NA 0.4; Olympus, Japan), 4x (UPlanFLN, NA 0.13; Olympus, Japan). The total magnification could be increased with an additional 1.6x magnification changer. Sperm, *Amoeba proteus*, and *Hydra vulgaris* were investigated in a custom-made observation chamber of 150 μm depth^39^. Images of *D. melanogaster* were recorded with a stereomicroscope (SZX12; Olympus, Japan), equipped with a DF PLAPO 1X PF lens (NA 0.11; Olympus, Japan).

### Characteristics of the multifocal adapter

The multi-channel imaging device QV-2 (Photometrics, USA) was used to split the light coming from the microscope into four light paths (Supplementary Fig. 1a). The light within each optical path was projected to a different location on the camera chip. The focal length of each optical path was varied by an additional lens located at the position designed for holding filters in the QV-2. For measurements, a set of lenses with the following focal lengths was used (in mm): *f*_1_ = ∞ (no lens), *f*_2_ = 1,000, *f*_3_ = 750, and *f*_4_ = 500 (Supplementary Fig. 1a).

### Intensity normalization across planes

Local differences of image intensity in the four focal planes (Supplementary Fig. 12a) were measured by recording an image without a specimen and generating an intensity heat-map (Supplementary Fig. 12b). For normalization, pixel intensity was divided by the respective fractional intensity value in the heat-map (Supplementary Fig. 12c).

### Alignment of plane images

The four focal planes were aligned at each time step using the ImageJ plugin *Multi-Stack Reg*.^71^. An alignment matrix was determined from the first time point and applied to all subsequent frames. Because the alignment algorithm uses the first plane as a template and corrects the other planes based on the first plane, this technique also allowed to correct for minute magnification differences appearing in planes 2-4 while not in plane 1 (Supplementary Table 1, Supplementary Table 2).

For the images of freely-swimming sperm, an image of a calibration grid was recorded (Supplementary Fig. 2a) and processed in ImageJ by image segmentation (threshold algorithm *Li*^72^, Supplementary Fig. 2b), skeletonization^73^ (Supplementary Fig. 2c), and alignment (*Multi-Stack Reg*^71^; Supplementary Fig. 2d). The output alignment matrix was used for plane alignment.

### Determination of the inter-plane distances

To determine the inter-plane distances, a series of multifocal images of a calibration grid at different positions set by a piezo was acquired. For each of the four planes, it was determined the piezo position at which a reference grid was in focus. The focus position was defined as the position at which the grid image featured the highest standard deviation of pixel intensity^74^. From the differences in focus position between neighboring planes, the inter-plane distances were determined.

For low magnification (stereomicroscope, 4x), where the piezo range was insufficient for spanning all planes, the inter-plane distance was estimated manually by measuring the required focus displacement between two adjacent planes for sharp imaging of the specimen in each plane.

### Simulating images produced by the multifocal adapter

Based on the inter-plane distances and the focal lengths of the lenses in the QV-2, we performed numerical calculations of images produced by the setup. The different imaging planes produced by the MFI setup were obtained with ray-optics calculations based on the thin-lens equation.

Defocusing was simulated by convolving the point spread function (PSF) with the sharpest image that we acquired of the calibration grid. We assumed a Gaussian PSF with transverse intensity distribution:

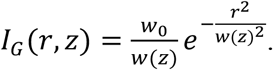

In the previous equation, 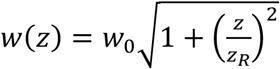 is the evolution of the Gaussian beam width with the propagation distance *z*, 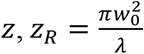 is the Rayleigh range, *w*_0_ is the Gaussian beam width at the focal plane, and *λ* is the wavelength. *w*_0_ is given by the optical resolution provided by the experimental set-up (Abbe limit, about 2.1 μm). The Abbe limit was calculated as 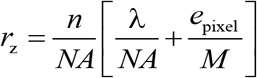, where *λ* is the wavelength, *n* is the refractive index, *NA* is the numerical aperture of the objective, *e*_pixel_ is the lateral pixel size, and *M* is the magnification of the objective. Based on the resolution, the inter-plane distances, and the Rayleigh range *z_R_*, we calculated the Gaussian beam width *w*(*z*) and PSF at every *z* and for each plane.

### Extended depth-of-field (EDOF)

Multifocal images were processed into extended depth-of-field (EDOF) images using the complex wavelet mode^24^ of the ImageJ plugin “extended depth-of-field” by “Biomedical Imaging Group”, EPFL, Lausanne. We created a customized version of this ImageJ plugin to enable automated processing of time series of image stacks.

### Recording Brownian motion

Carboxylate-modified latex beads of 0.5 μm diameter (C37481, Lot # 1841924, Thermo Fisher Scientific Inc., USA) in an aqueous solution were inserted into the custom-made observation chamber. Images were recorded with the 20x objective (NA 0.5) and the additional 1.6x magnification changer at 500 fps. A 530-nm LED was used as a light source. For each timepoint, a maximum-intensity-projection across the four planes was generated. Bead positions were analyzed as described in the next chapter.

### Tracking beads in multifocal images

For each time-point, a maximum-intensity-projection across the four planes was generated. To remove non-moving beads and background structures from images of beads around sperm, a time-average projection was subtracted from all time-points. To determine bead positions in x, y, and time, the maximum-intensity-projection time series was subjected to the FIJI plugin “TrackMate”^75^ (settings: LoG detector, estimated blob diameter of 10 pixel, threshold of 50, sub-pixel localization).

Each bead’s z-position was analyzed using a java-based custom-written ImageJ plugin using the “TrackMate”-derived xy and time coordinates as a template.

For each of the focal planes, all pixels within a radius of 10 px (about 3.4 μm) were subjected to an unconstraint 2D radial fit to retrieve the radius. The fit was developed as follows.

For a circle of radius *R* centered at *C* = (*C*_0_, *C*_1_), the circle equation (*x* – *C*_0_)^2^ + (*y* – *C*_1_)^2^ = *R*^2^ can be rewritten as (2*x*)*C*_0_ + (2*y*)*C*_1_ + *C*_2_ = *x*^2^ + *y*^2^, where 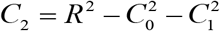. With the coordinates of the pixel given, we can fit the transformed circle equation linearly to calculate the parameters *C*_0_, *C*_1_, and *C*_2_. The linear fit is constructed into the matrix form **A***c* = **B**, where 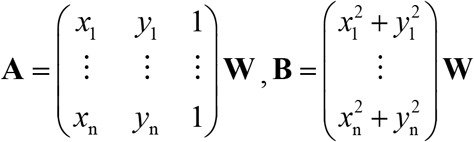, and 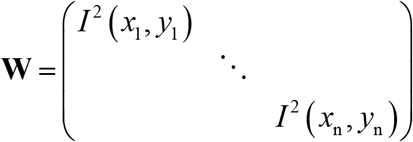 is the weight matrix defined by the square of the pixel intensity. 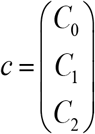 is the approximate solution of this over-determined equation, which can be calculated with least-squares principle. Finally, the circle radius *R* can be determined using the equation: 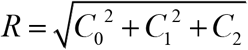.

If the quality of the fit *r*^2^> 0.8, the radius was used to infer the z-position. If the fit to the radial profile did not match the criterion in more than two planes, the bead was excluded from analysis and no z-position was determined. The z-position was determined for each remaining bead as follows. In each focal plane, a difference function of the determined bead radius and the plane’s calibrated relationship between bead radius and z-position was determined. Each difference function shows two minima, which represent the two most probable z-positions (one located above and another below the focal plane). Finally, the position of the bead was unequivocally determined from the multiple measures of the z-positions across different planes: the combination of z-positions from the different planes with the lowest standard deviation was selected. For each z-position, the precision was predicted based on the calibrated relationship of z-position and fit width. The z-position with lowest precision value was selected as the final bead’s z-position.

Images of immobilized beads at different heights set by a piezo-driven objective were used for calibration of z-positions. Radius profiles across z of seven different beads were aligned by minimizing the least-mean-square between all the planes. Finally, aligned profiles were averaged and smoothed. This resulted in a radius function R(z). To estimate the precision error σ_z_ resulting from inferring the z position of beads using this function, we used error propagation of the inverse function: 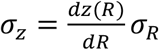, where σ_R_ is the standard deviation of the calibration for different beads.

Individual bead coordinates on each frame were connected to form tracks using custom-made software written in MATLAB. The software searched for the nearest particle across frames within a distance of less than 1.5 μm.

### Determining the x-,y-, and z-precision based on Brownian motion of particles

For characterizing Brownian motion, the normalized histogram of bead displacement along each dimension *d* (*d* = *x, y*, or *z*) of space was fitted to the characteristic probability density function of diffusive particles^76^:

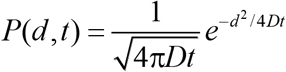

The measured distribution of bead displacement Δ*s* results from positional variations due to diffusion during each time interval Δ*t* superimposed to positional variations that result from measurement errors due to the precision of the multifocal method. Assuming that measurement errors are independent of diffusion, the distribution of bead displacement measured *P*_M_(Δ*s, t*) can be described by the convolution of the two probabilistic distributions characterizing the measurement errors due to precision *P*_p_(Δs) and diffusion *P*_D_(Δ*s*, Δ*t*):

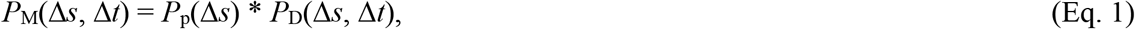

where * denotes the convolution operation. Assuming isotropic diffusion and normally distributed measurement errors, it can be readily found (use the convolution theorem for this) that the variance of the resulting distribution σ_M_^2^ is characterized by the sum of the variances:

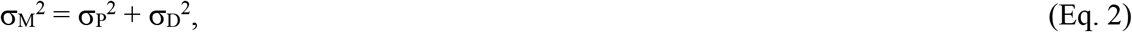

where the standard deviation σ_p_ characterizes the precision of the method, and σ_D_^2^ the variance of the bead displacement due to diffusion. For each dimension of space, the corresponding diffusive variance can be calculated provided that the diffusion coefficient *D* is known:

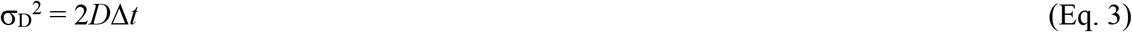

For a spherical particle at high Reynolds numbers, the diffusion coefficient is well described by the Stokes-Einstein relation:

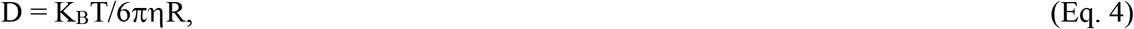

where *K*_B_ is the Boltzmann constant, *T* the temperature (295 K), *R* the particle radius (250 nm), and η(*T*) the dynamic viscosity of the embedding fluid (ca 0.95·10^-3^ kg·m^-1^·s^-1^ for water at 295 K). Combining Eqs. 2–4, we find that the precision of the method to be:

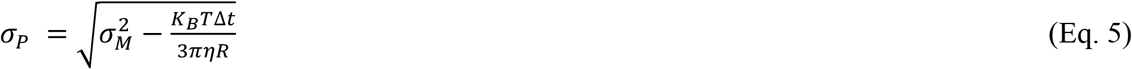

Using Eq. 5 with a frame period Δ*t* of 2 ms, we obtain a z precision of 150 nm (N = 82 beads recorded for a mean duration of 3.3 s), which is in agreement with the precision found during the calibration procedure (Fig. 2g). Additionally, the precision in the XY plane can be estimated by this procedure to be 45 and 37 nm for the x and y axis, respectively.

### Reconstruction of the flagellar beat

To reconstruct the flagellar beat, we developed a fully automated ImageJ plugin written in Java (SpermQ-MF). This plugin was based on SpermQ, a preexisting plugin for 2D analysis of flagellar beating^39^. The sharpest (highest standard deviation of pixel intensity) of the four plane images was subjected to the standard SpermQ workflow to reconstruct the flagellum in 2D. Next, the z-position of each point was determined using two methods based on the flagellar width or the pixel intensity.

For the method based on the width, a Gaussian curve was fitted to the image intensity in the direction normal to the flagellum at that particular point. The width of the resulting Gaussian was then converted into two z-positions (one located above and another below the focal plane) using the calibrated relationship between the flagellar width, the position on the flagellum, and the z-distance to the respective plane. Finally, the position of the flagellar point was unequivocally determined from the multiple measures of the z-positions across different planes.

For the z-localization method based on the changes of pixel intensity, we determined the intensity value of a given flagellar point in each plane by linear interpolation from the four surrounding pixels weighted by their distance to the given point. Next, each intensity value, together with the focal position of the corresponding plane, formed a point in a coordinate system of focal position and intensity. A Gaussian curve was fitted to the resulting four points, whose center referred to the z-position. All determined z-positions were averaged and smoothed along the arc length as described^39^.

z-positions were calibrated using recordings of non-motile sperm at different heights set by a piezo-driven objective. Calibrations from all different analyzed sperm (human: *n* = 12 sperm from five different donors, sea urchin: *n* = 5 sperm) were averaged and smoothed by a 6×6 kernel median-filter.

For each sperm, the x-,y-,z-, and time coordinates of the flagellar points were further analyzed in MATLAB to plot the flagellar trajectories and to determine the beating plane, flagellar curvatures, and frequency spectra. In 3D visualizations of the flagellum, only arc lengths 5 – 50 or 5 – 45 μm were shown for human or sea urchin sperm, respectively.

For determining the flagellar beat plane, Gyration Tensor analysis was performed^29^. The rolling speed was determined based on the normal vector of the beat plane. The rolling frequency was calculated from the simple average of the rolling speed. The curvature was calculated with a moving window (4 μm arc length) along the flagella. Within the moving window, the flagellar segment was projected to the osculating plane and Taubin’s method^77^ was used to fit the osculating circle of the projected flagellar segment. This algorithm is available at: https://de.mathworks.com/matlabcentral/fileexchange/47885-frenet_robust-zip.

For each sperm, the frequency spectrum was obtained by subjecting the time-course of the 3D curvature at arc lengths 33 μm (human sperm) or 34.4 μm (sea urchin sperm) to the MATLAB function “pwelch”, determining Welch’s power spectral density, with the following settings (moving window of 200 pixels, moving step of 40 pixels, 1 pixel = 11/32 μm). The *C*_0_, *C*_1_, and *C*_2_ values were estimated by fast Fourier transform of flagellar curvature at arc lengths 33 μm (human sperm) or 34.4 μm (sea urchin sperm).

### Flow in aqueous solution around a human sperm cell

Sperm were suspended in an HTF solution containing 100 μg/ml HSA (Ref. 9988, Irvine Scientific, USA) and 40 μg/ml carboxylate-modified latex beads of 0.5 μm diameter (C37481, Lot # 1841924, Thermo Fisher Scientific Inc., USA), and inserted into the custom-made observation chamber. Images were recorded with the 20x objective (NA 0.5) and the additional 1.6x magnification changer at 500 fps. The flow field was calculated from the tracked bead positions using a custom-made program written in MATLAB. The volume was divided in a mesh grid xyz of volume 4 x 4 x 2 μm. The spatial and temporal mean velocity of particles within that volume was calculated to produce an estimate of the flow velocity within each voxel. The z glass coordinate was estimated as the mean z-position of 114 non-moving beads.

### Swimming trajectories and swimming speed

Sperm swimming trajectories were smoothed by Matlab in-built smooth function with the “rlowess” method and a time window of 8 ms. The centreline of sperm swimming trajectories was smoothed with the “rlowess” method and a time window of 80 ms. The swimming speed was estimated by the distance of two points along the centreline divided by the time interval.

### Software

Image processing and analysis were performed in ImageJ (v1.52i, National Institutes of Health, Bethesda, MN, USA) and MATLAB 2018b (Mathworks). Calculations were performed in MATLAB 2018b (Mathworks), Rstudio (version 1.2.5019), and R (version 3.6.1). Plots and figures were generated using GraphPad Prism (Version 6.07, GraphPad Software, Inc., La Jolla, CA, USA), OriginPro (Version 9.0.0G, OriginLab Corporation, Northampton, USA), MATLAB 2018b (Mathworks), and Adobe Illustrator CS5 (Adobe Systems, Inc., v15.0.0, San Jose, CA, USA). The Java-based software was developed and compiled using Eclipse Mars.2 (Release 4.5.2, IDE for Java Developers, Eclipse Foundation, Inc., Ottawa, Ontario, Canada) and using parts of the source code of SpermQ (v0.1.7,^39^).

## Data availability

Exemplary high-speed video recordings to run the analysis routines, extracted data, and analysis supporting this study are freely accessible via the GitHub repository https://github.com/hansenjn/MultifocalImaging-AnalysisToolbox. Additional data that support the findings of this study are available from the corresponding author upon request.

## Code and software availability

The developed ImageJ plugins along with the source code, manual, and example data are freely accessible on GitHub (https://github.com/hansenjn/MultifocalImaging-AnalysisToolbox). Like ImageJ, all the developed ImageJ plugins are open-source.

## Acknowledgement

This work was funded by the German Research Foundation (DFG) under Germany’s Excellence Strategy – EXC2151 – project number 390873048 (D.W.), SPP1726 “Microswimmers” (L.A., U.B.K, J.F.J, D.W.), as well as the Boehringer Ingelheim Fonds (J.N.H.) and the Alexander von Humboldt Foundation (A.T.).

## Author Contribution

J.F.J, L.A., D.W., and U.B.K. conceived the project. J.N.H., J.F.J., L.A., and U.B.K. designed the experiments. J.N.H., J.F.J., A.G., and R.P. performed the experiments. J.N.H., L.A., J.F.J., and A.G. developed software to analyze the data. J.N.H., A.G., J.F.J., and L.A. analyzed the data. A.T. performed theoretical estimations. J.N.H., L.A., J.F.J., and U.B.K. wrote the manuscript. All authors commented on and edited the text.

