## Supplementary Information for "Multifocal imaging for precise, label-free tracking of fast biological processes in 3D"

#### Supplementary Figures

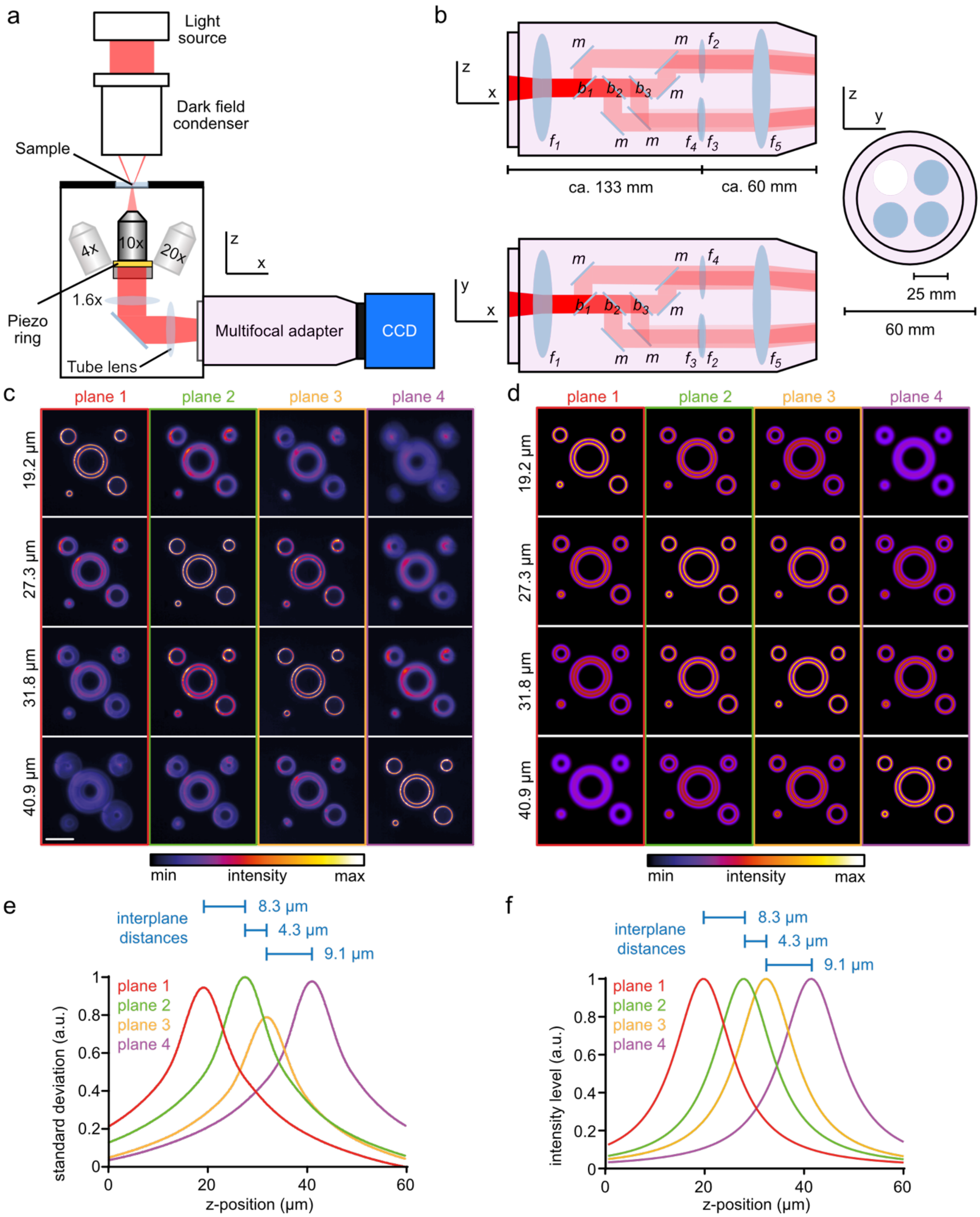


**Supplementary Figure 1 | The multifocal adapter. (a)** Composition of the simple multifocal imaging setup. The multifocal adapter (MFA) is inserted into the light path between a standard dark-field microscope and a camera. **(b)** Composition of the multifocal adapter. Incoming light from the microscope is infinity projected by the lens $f_1$ and split by three consecutive beamsplitters ($b_1$ = 25/75; $b_2$ = 33/67; $b_3$ = 50/50). Five mirrors (*m*) guide the light into four separate optical paths. The lens $f_5$ projects the four images onto the camera chip. The inter-plane distance is set by lenses of different focal length ($f_1$ - $f_4$). **(c, d)** Experimentally acquired (c) and simulated (d) (see Materials and Methods) multifocal dark-field images (planes 1 - 4, 20x objective) of a calibration grid at the four focal positions (indicated on the left side), where one of the four focal planes maps the grid sharply. Scale bar represents 20 µm. **(e, f)** Experimentally determined (e) and simulated (f) standard deviation of pixel intensity (values linearly rescaled for each plane) as a function of the piezo-controlled objective z-position (0.1 µm steps; 20x objective). The z-position corresponding to the sharpest image in each plane was determined as the position where the normalized standard deviation of the image was maximal. Blue bars represent the interplane distances.

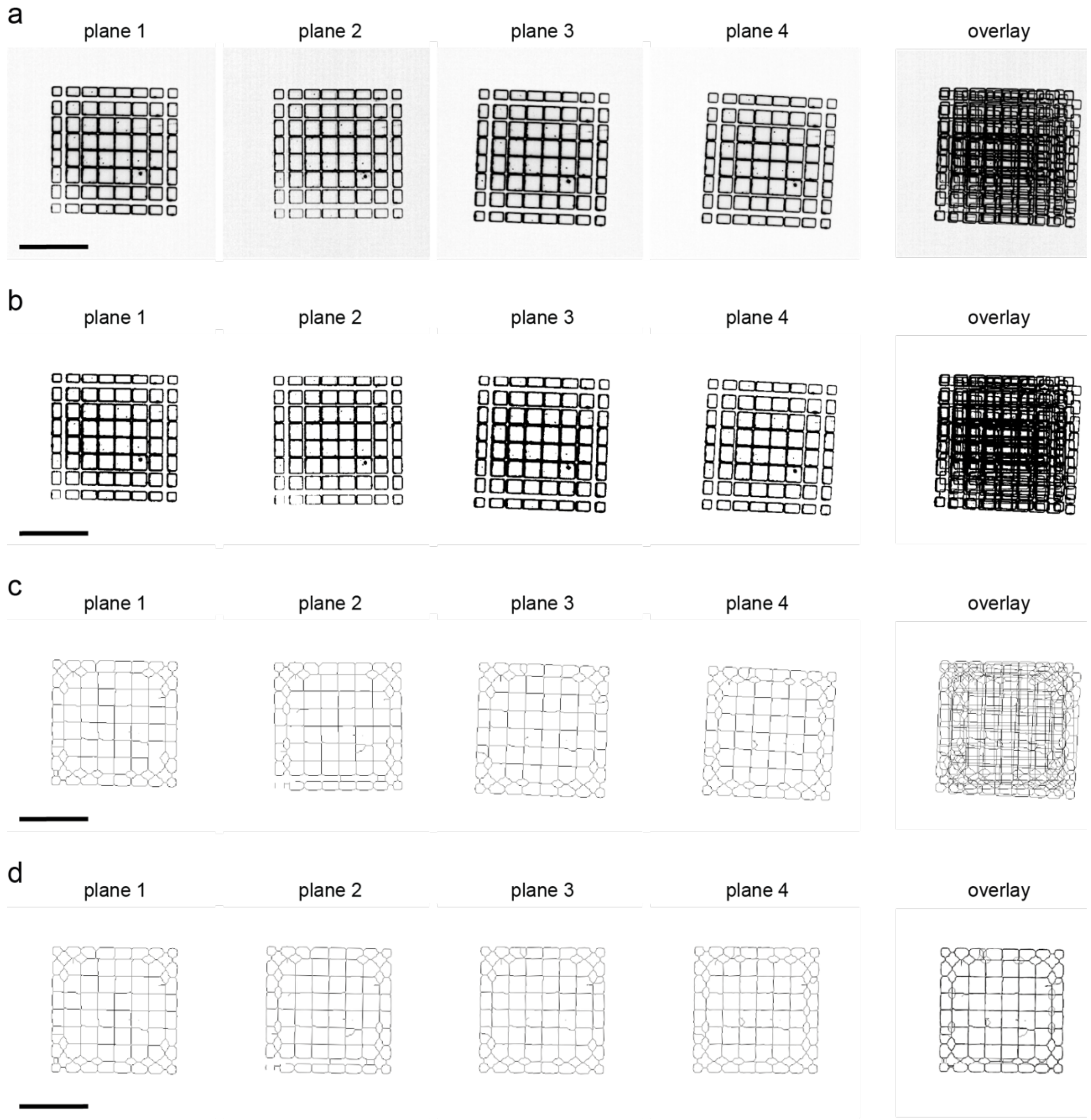


**Supplementary Figure 2 | Alignment of plane images in multifocal recordings. (a)** A multifocal image of a calibration grid was recorded to generate an alignment matrix for the four plane images. **(b)** The image was segmented via ImageJ's thresholding algorithm *Li* [1] and **(c)** subsequently skeletonized via the FIJI plugin “Skeletonize3D” [2]. **(d)** The skeleton images were aligned by the ImageJ plugin “Multi-Stack Reg” [3], and the registration matrix was stored for alignment of all data acquired with the same magnification. Image intensities were inverted for better visualization. Scale bars represent 100 µm.

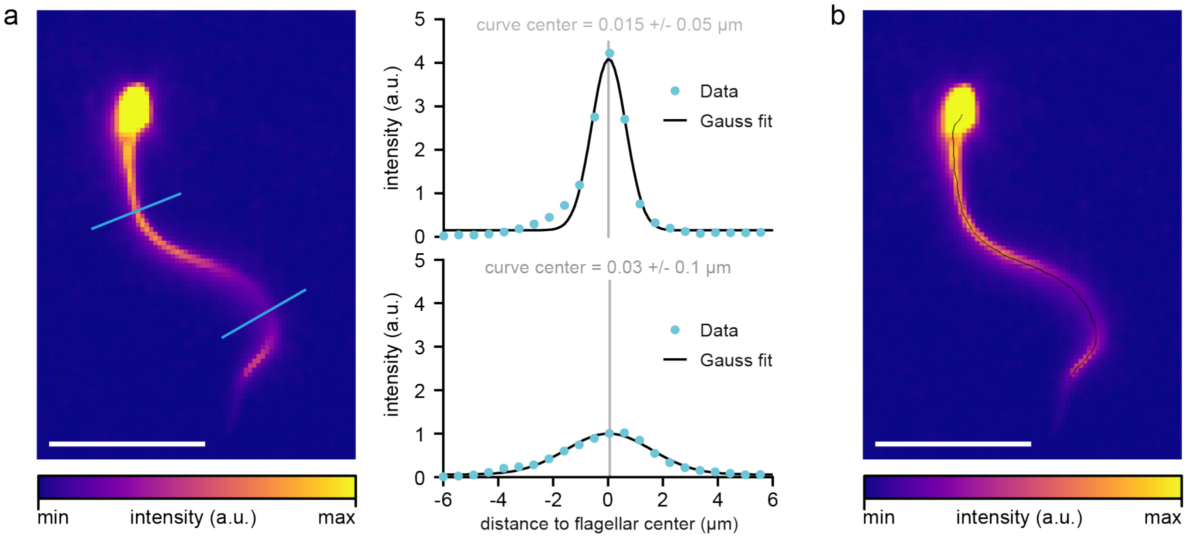


**Supplementary Figure 3 | High-precision reconstruction of the flagellum in 2D. (a)** Exemplary plane image from a multifocal recording of a human sperm cell. To reconstruct the x and y coordinates of the flagellum, the image was analyzed with SpermQ [4]. SpermQ first generates an initial coarse skeleton and then fits the intensity profile on normal lines along the flagellum to Gaussian curves. Two exemplary normal lines are shown in cyan and their corresponding intensity profiles with Gaussian curve fits on the right. SpermQ corrects the flagellar reconstruction by shifting the reconstructed points to the Gaussian-curve centers (indicated in grey, including 95% confidence intervals). Of note, the error of the Gaussian curve fits reveals a high precision in x and y (0.05 µm, top, and 0.1 µm, bottom). **(b)** This adjustment procedure allows determining flagellar coordinates (black line) with high precision. Scale bars represent 20 µm.

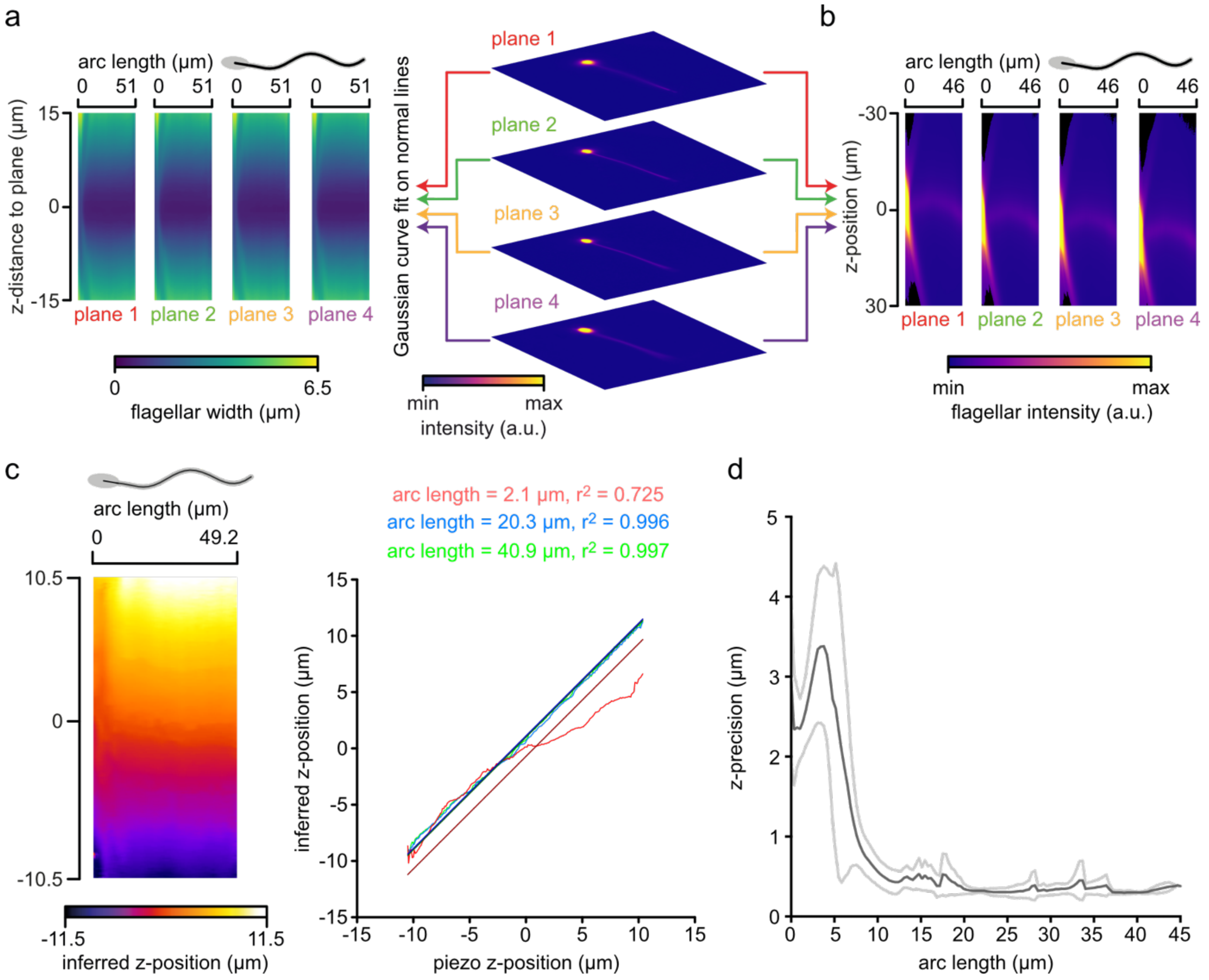


**Supplementary Figure 4 | Reconstructing flagellar z-positions of sea urchin sperm using four focal planes.**

**(a-b)** Characterizing the relationship between image and z-position of sea urchin sperm flagella in the MFI setup. **(a)** Flagellar width (color-coded), determined by a Gaussian curve fit on a normal to the flagellum, as a function of the flagellar position (arc length) and the z-distance to the respective plane (mean image of $n$ = 5 sperm). **(b)** Maximum flagellar intensity (color-coded) as a function of the flagellar position (arc length) and the z-position relative to the four planes. **(c)** z-position (color-coded) along the flagellum (arc length) of an immotile sea urchin sperm inferred from MF images based on the calibrated relationship between the flagellar width, position on the flagellum, and z-distance to the respective plane during modulation of the objective z-position with a piezo (steps 0.1 µm; left). Three different flagellar positions are further analyzed by a linear curve fit with a slope constraint to unity (right). **(d)** Standard deviation (SD) of the residuals of linear curve fits to data as exemplified in (c). Mean ± standard deviation of $n = 3$ sperm.

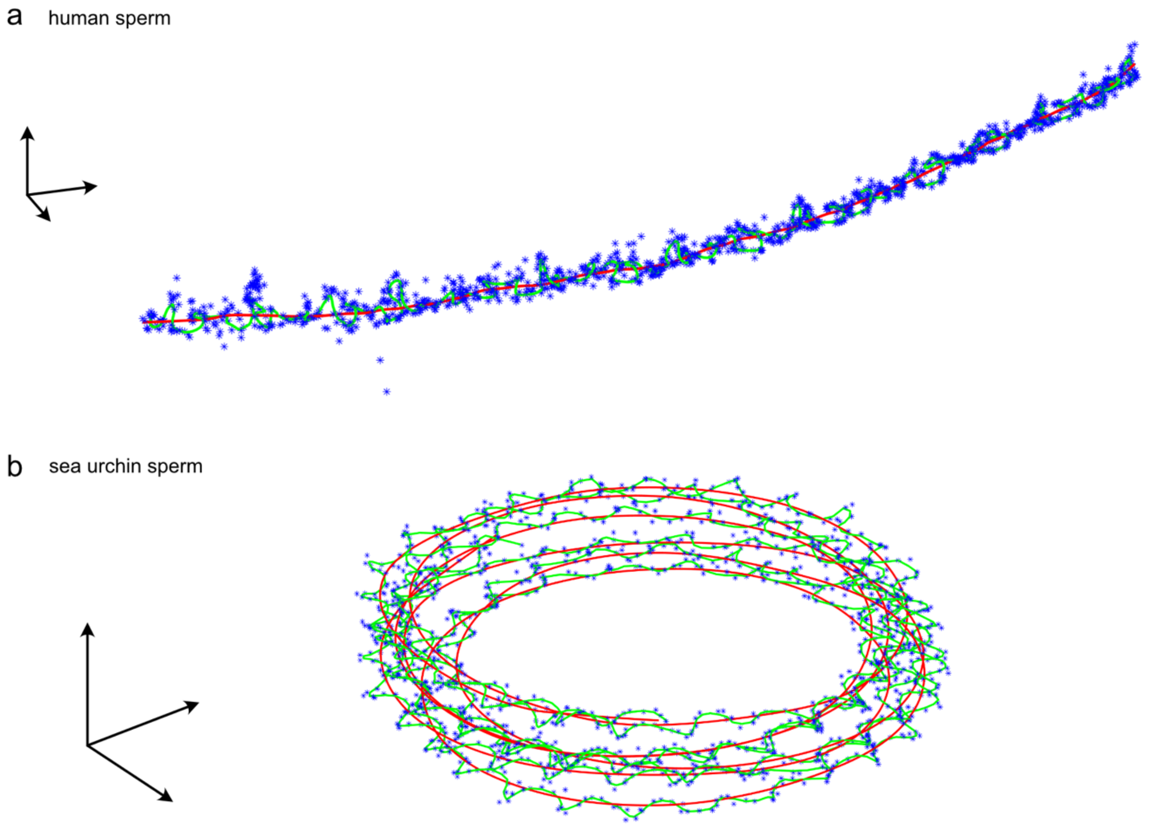


**Supplementary Figure 5 | Determining the swimming speed of freely swimming human and sea urchin sperm.** Tracking of the head centre from freely swimming **(a)** human and **(b)** sea urchin sperm. Tracked positions (blue asterisks) and smoothed head position with smoothing time windows of 20 ms (green line) and 200 ms (red line) are shown. The red line is used to determine the swimming speed. Arrows: 20 µm.

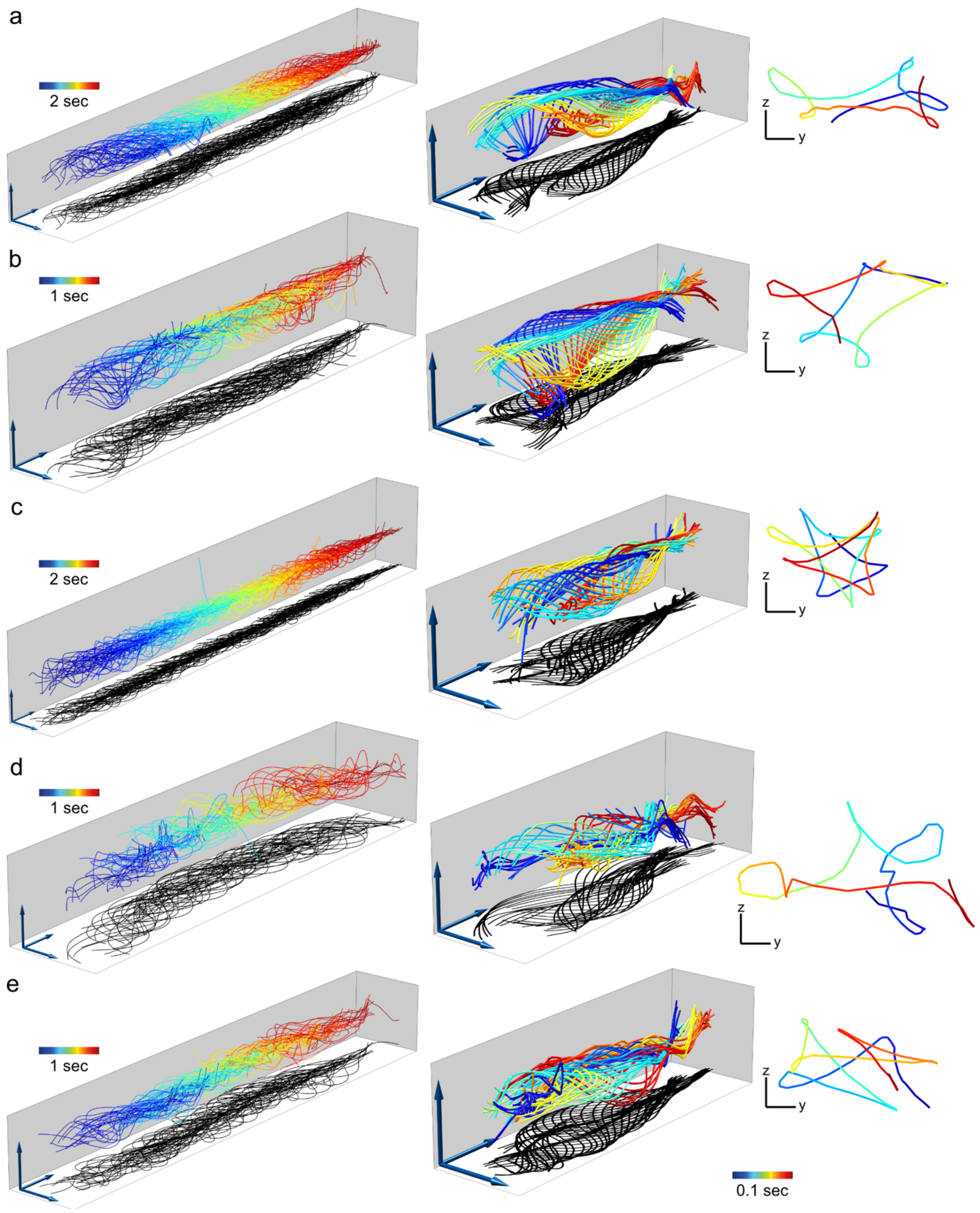


**Supplementary Figure 6 | Trajectories and beat patterns of free-swimming human sperm.** 3D-tracked flagella from exemplary free-swimming human sperm. Panels **(a-e)** represent individual tracked sperm cells. Views of the whole timespan tracked to show the trajectory (left, only every 4$^{th}$ frame plotted for better visualization), of a selected timespan for better visualization of the flagellar beat (middle), or of a Y-Z-projection of a selected flagellar point (arc length 20 μm, view in swimming direction) for visualization of sperm rolling. Bars and arrows indicate 10 μm.

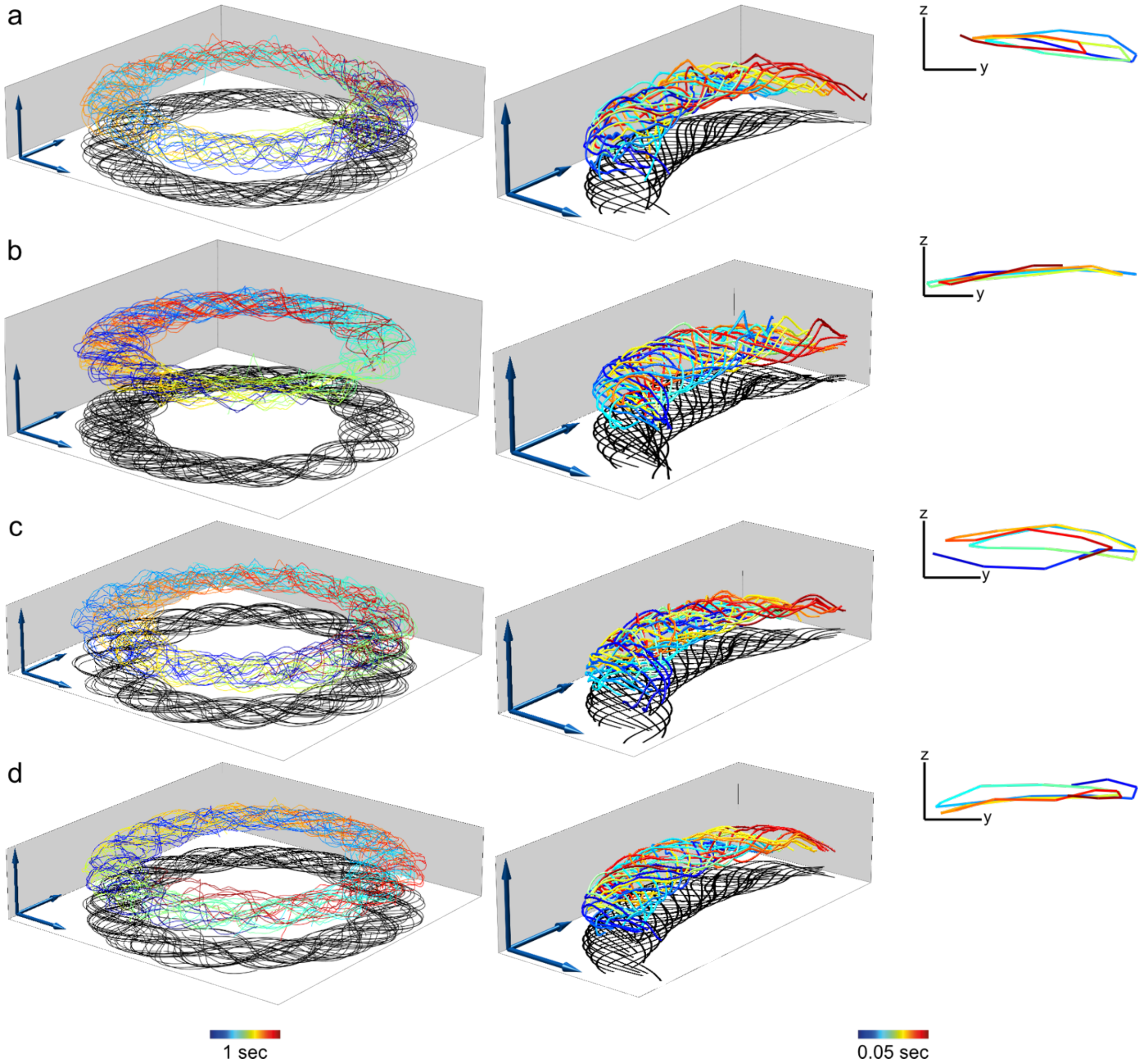


**Supplementary Figure 7 | Trajectories and beat patterns of free-swimming sea urchin sperm.**

3D-tracked flagella from exemplary free-swimming sea urchin sperm. Panels **(a-d)** represent individual tracked sperm cells. Views of the whole timespan tracked to show the trajectory (left, only every 4th frame plotted for better visualization), of a selected timespan for better visualization of the flagellar beat (middle) or of a Y-Z-projection of a selected flagellar point (arc length 20 µm, view in swimming direction) for visualization of sperm rolling. Bars and arrows indicate 10 µm.

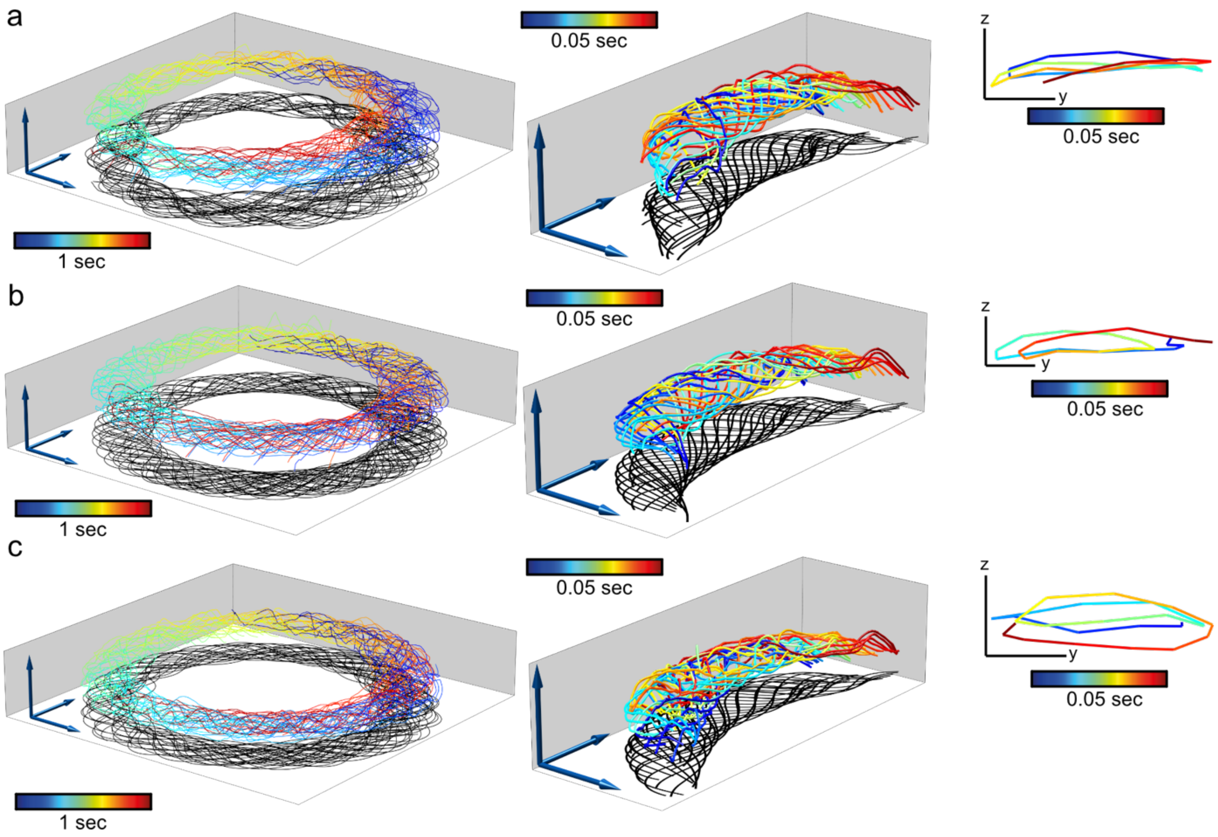


**Supplementary Figure 8 | Trajectories and beat patterns of free-swimming sea urchin sperm.** Continuation of Supplementary Figure 7.

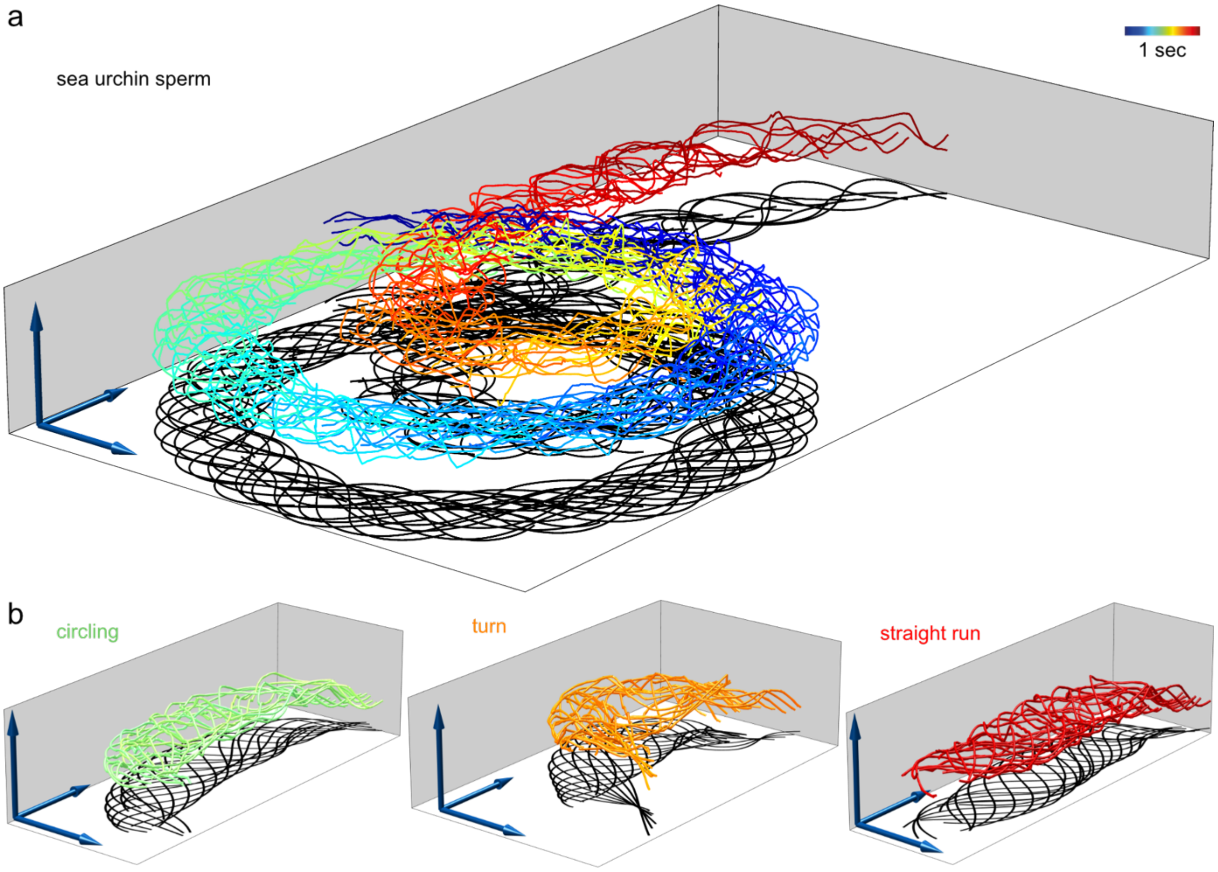


**Supplementary Figure 9 | Trajectories and beat patterns of a sea urchin sperm during a pronounced change in swimming behavior.** 3D-tracked flagella from a freely swimming, sea urchin sperm. **(a)** Projection of 1 s of flagellar beating (for better visualization only every 4$^{th}$ frame was plotted). **(b)** Projection of selected time periods (0.05 sec) during swimming on a circle (left), while looping (middle), and swimming straight (right). Projections in (b) are color-coded according to the time-colour-map in (a). Arrows indicate 10 µm.

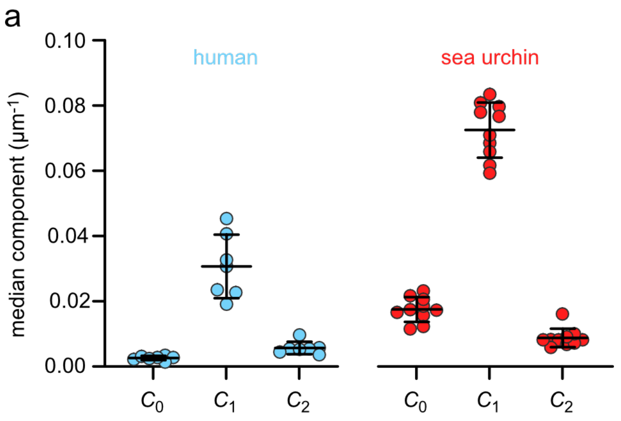


**Supplementary Figure 10 | Extended analysis of the flagellar beat of free-swimming human and sea urchin sperm. (a)** Decomposition of the flagellar beat wave into three curvature components: a time-average bending of the flagellum $C_0$, the curvature amplitude $C_1$ at the fundamental beat frequency, and the second harmonic $C_2$. Each data-point corresponds to the median for an individual sperm.

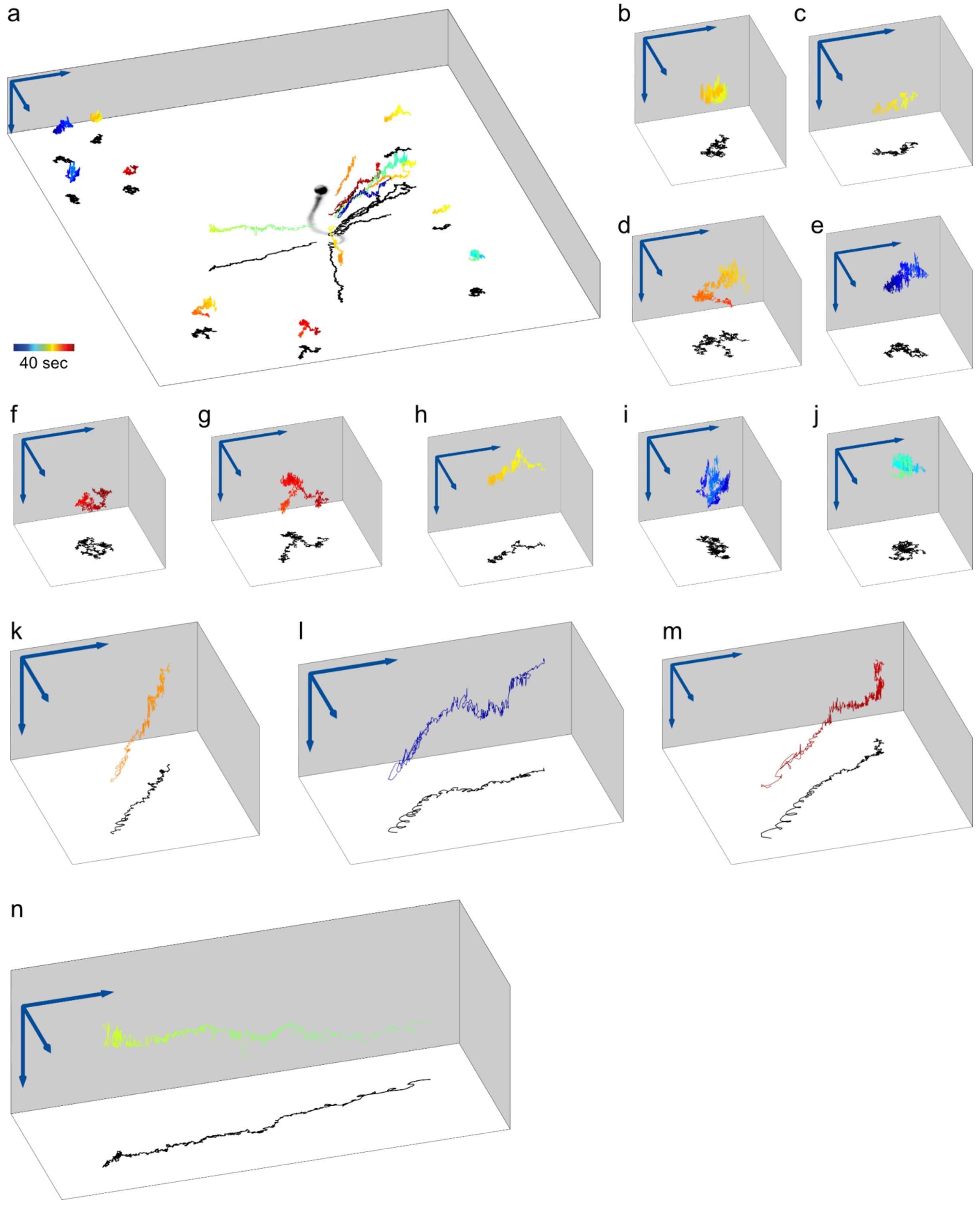


**Supplementary Figure 11 | Exemplary trajectories of latex beads around a human sperm cell.** Additional magnified views from Figure 6c. **(a)** Trajectories of few exemplary beads from Fig. 6b (image copied from Figure 6c). Time color-coded as indicated. Individual trajectories magnified in (b) to (n). **(b-j)** Trajectories of beads remote from the flagellum-driven fluid flow.

**(k-m)** Trajectories of beads that are attracted to the sperm cell. **(n)** Trajectory of a bead that is moving away from the flagellum.

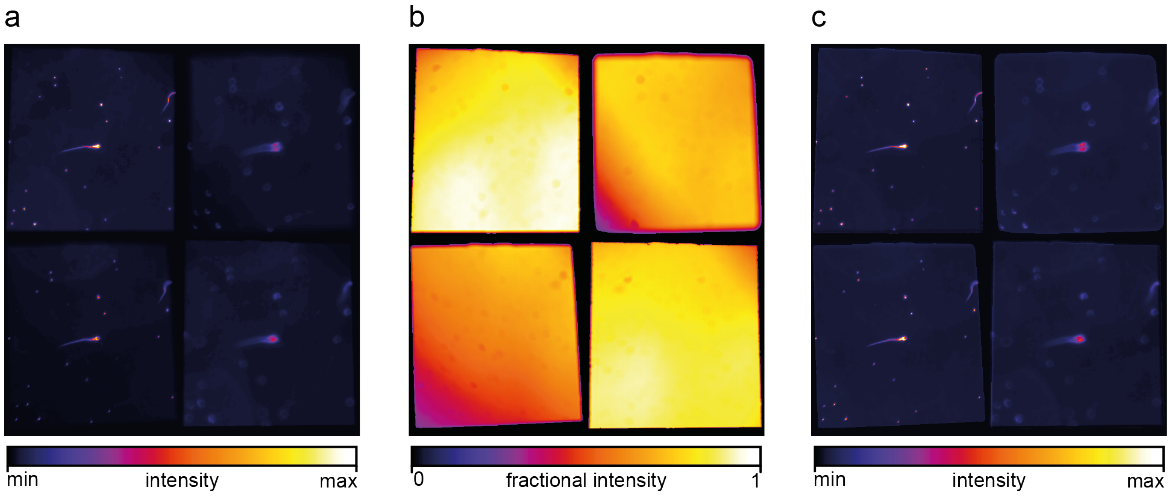


**Supplementary Figure 12 | Local intensity correction of images. (a)** Multifocal image of a human sperm cell (20x objective). **(b)** Normalized intensity heat-map recorded for the MFI set-up by acquiring images without a specimen. **(c)** Intensity-corrected image after dividing each pixel intensity by the respective fractional intensity value in the intensity heat-map (b).

### Supplementary Tables

**Supplementary Table 1 | Predicted and measured pixel size in the different plane images of the uncorrected multifocal images for a set of lenses[1]**

| Magni-fication | Objective | Magnification changer | Pixel size (µm / px) in x / y | | | | |
|---|---|---|---|---|---|---|---|
| | | | Predicted | Plane 1 | Plane 2 | Plane 3 | Plane 4 |
| 4x | 4x, NA 0.13 | None | 2.75 | 2.74 / 2.75 | 2.80 / 2.79 | 2.80 / 2.80 | 2.80 / 2.86 |
| 6.4x | 4x, NA 0.13 | 1.6x | 1.72 | 1.72 / 1.73 | 1.74 / 1.73 | 1.73 / 1.73 | 1.76 / 1.75 |
| 10x | 10x, NA 0.4 | None | 1.10 | 1.10 / 1.10 | 1.12 / 1.12 | 1.13 / 1.13 | 1.16 / 1.15 |
| 16x | 10x, NA 0.4 | 1.6x | 0.69 | 0.69 / 0.70 | 0.70 / 0.70 | 0.70 / 0.70 | 0.71 / 0.70 |
| 20x | 20x, NA 0.5 | None | 0.55 | 0.55 / 0.55 | 0.56 / 0.56 | 0.56 / 0.56 | 0.57 / 0.57 |
| 32x | 20x, NA 0.5 | 1.6x | 0.34 | 0.35 / 0.34 | 0.35 / 0.35 | 0.35 / 0.35 | 0.35 / 0.35 |

[1] in mm: $f_1=\infty$ (no lens), $f_2 = 1000$, $f_3 = 750$, and $f_4 = 500$

**Supplementary Table 2 | Predicted and measured pixel size in the multifocal images after alignment of plane images (Supplementary Fig. 2) for a set of lenses[1]**

| Magnification | Objective | Magnification changer | Pixel size (µm / px) | | |
|---|---|---|---|---|---|
| | | | Predicted | Measured in x | Measured in y |
| 4x | 4x, NA 0.13 | None | 2.750 | 2.747 | 2.749 |
| 6.4x | 4x, NA 0.13 | 1.6x | 1.719 | 1.722 | 1.732 |
| 10x | 10x, NA 0.4 | None | 1.100 | 1.100 | 1.101 |
| 16x | 10x, NA 0.4 | 1.6x | 0.688 | 0.692 | 0.695 |
| 20x | 20x, NA 0.5 | None | 0.550 | 0.547 | 0.548 |
| 32x | 20x, NA 0.5 | 1.6x | 0.344 | 0.345 | 0.344 |

[1] in mm: $f_1=\infty$ (no lens), $f_2 = 1000$, $f_3 = 750$, and $f_4 = 500$

**Supplementary Table 3 | Distances of planes 2, 3, and 4 to plane 1 for a set of lenses[1]**

| Magnification | Objective | Magnification changer | Distance (µm) | | |
|---|---|---|---|---|---|
| | | | Plane 2 | Plane 3 | Plane 4 |

| | | | | | |
|---|---|---|---|---|---|
| 4x | 4x, NA 0.13 | None | 231 | 363 | 573 |
| 6.4x | 4x, NA 0.13 | 1.6x | 45 | 67.2 | 113.6 |
| 10x | 10x, NA 0.4 | None | 36.8 | 54 | 90.2 |
| 16x | 10x, NA 0.4 | 1.6x | 14.6 | 21.6 | 35 |
| 20x | 20x, NA 0.5 | None | 8.3 | 12.6 | 21.7 |
| 32x | 20x, NA 0.5 | 1.6x | 3.5 | 4.7 | 8.8 |

[1] in mm: $f_1=\infty$ (no lens), $f_2$ = 1000, $f_3$ = 750, and $f_4$ = 500

### Supplementary Movie

#### Supplementary Movie 1

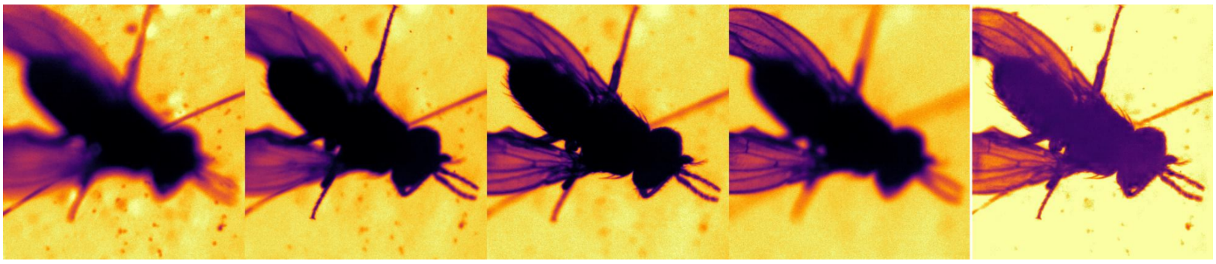

Multifocal imaging combined with an EDOF algorithm to visualize grooming *D. melanogaster* (1x magnification, play speed is real time).

#### Supplementary Movie 2

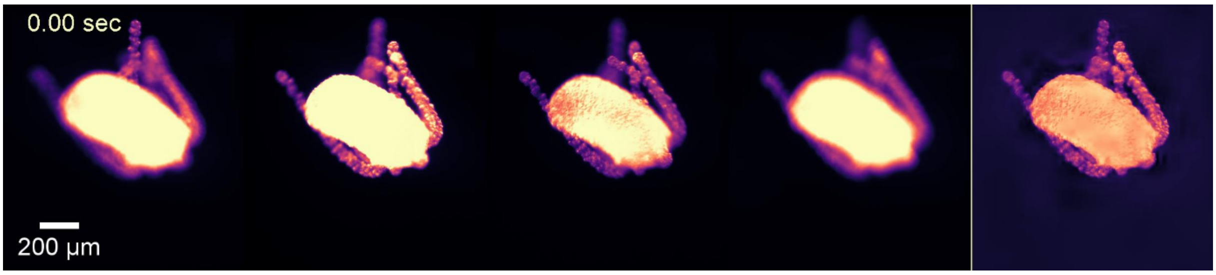


Multifocal imaging combined with an EDOF algorithm to visualize a foraging *Hydra vulgaris* (4x magnification).

#### Supplementary Movie 3

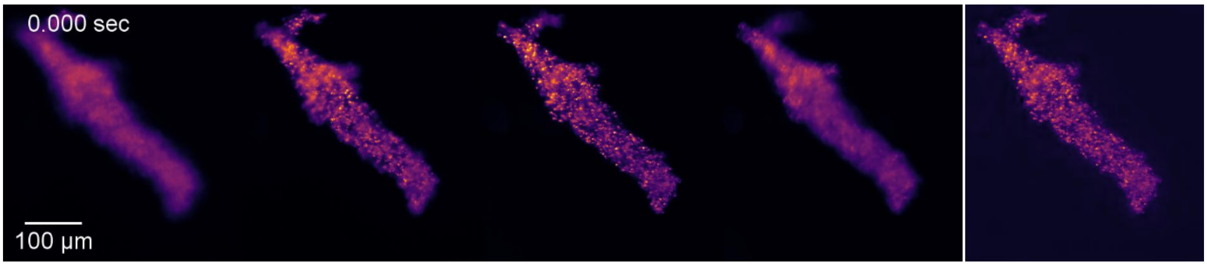


Multifocal imaging combined with an EDOF algorithm to visualize a crawling *Amoeba proteus* (10x magnification).

**Supplementary Movie 4**

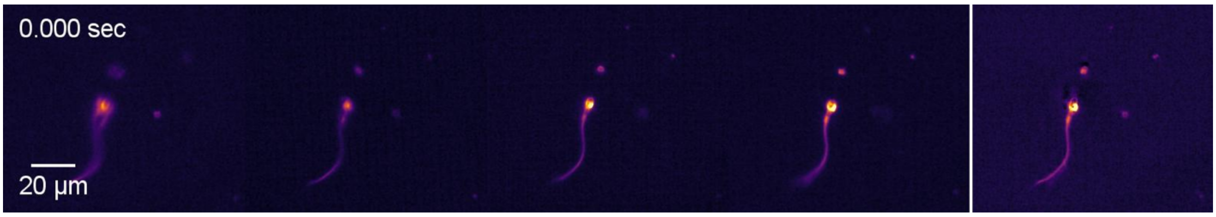


Multifocal imaging combined with an EDOF algorithm to visualize a swimming human sperm cell (32x magnification).

**Supplementary Movie 5**

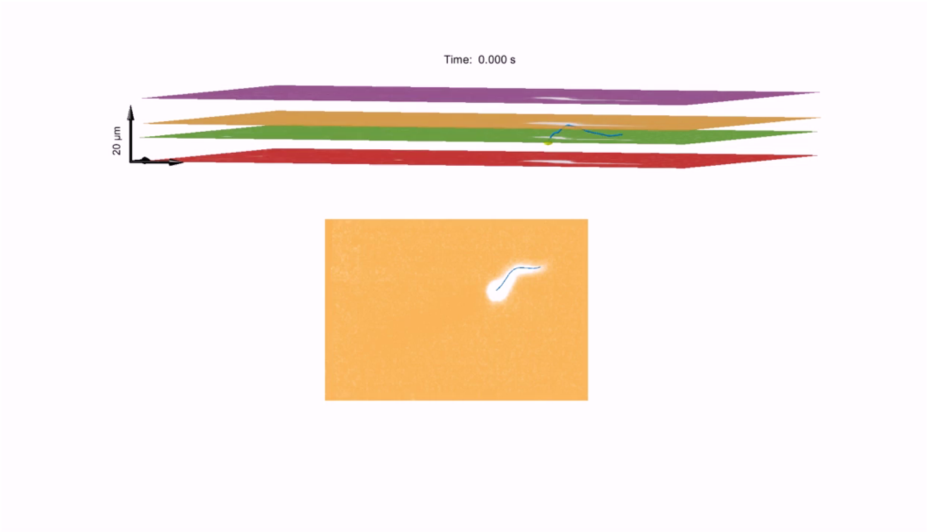


Top: 3D visualization of the four planes acquired by MFI and the flagellum reconstructed using SpermQ-MF and the calibrated relationship between flagellar width, position on the flagellum, and z-distance to the respective plane (Fig. 3a). Flagella indicated in blue. Positions of sperm heads indicated as yellow spheres. Bottom: Individual plane image acquired by MFI and the flagellum reconstructed in 2D (blue).

**Supplementary Movie 6**

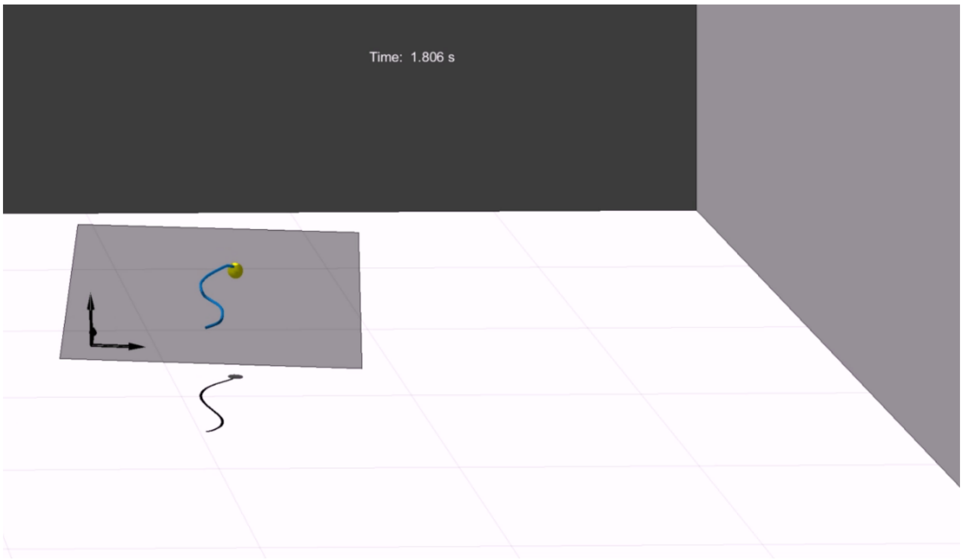


Flagellar 3D reconstruction of a free-swimming human sperm cell over time based on MFI and SpermQ-MF. The beat plane (gray) was defined by the Eigen-vectors of the flagellum.

**Supplementary Movie 7**

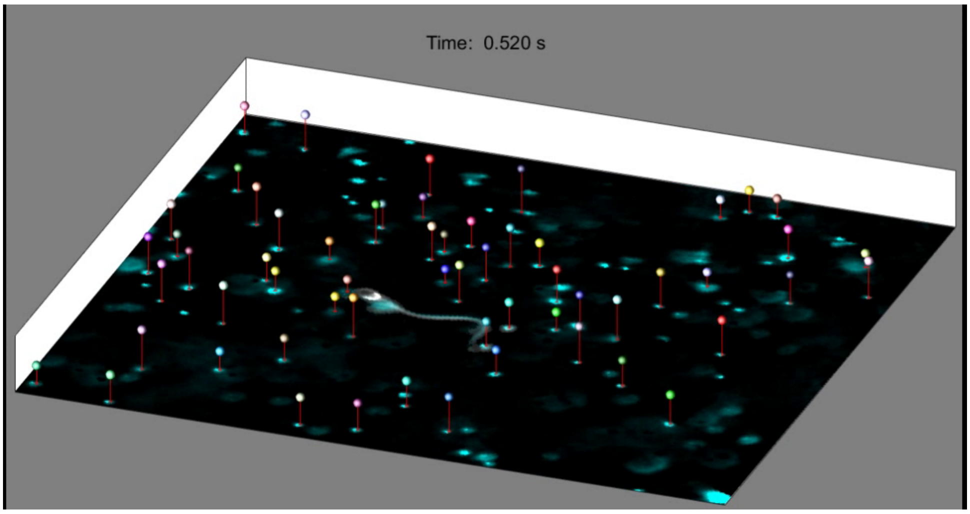


Tracking of latex beads flowing around a human sperm cell that was tethered to the cover glass at the head (each trajectory is depicted by a different colour). On bottom: selected plane image from the multifocal recording.
